# Urbanization and fragmentation interact to drive mutualism breakdown and the rise of unstable pathogenic communities in forest soil

**DOI:** 10.1101/2023.05.16.540503

**Authors:** Chikae Tatsumi, Kathryn F. Atherton, Sarah Garvey, Emma Conrad-Rooney, Luca L. Morreale, Lucy R. Hutyra, Pamela H. Templer, Jennifer M. Bhatnagar

## Abstract

Temperate forests are particularly threatened by urbanization and fragmentation, with over 20% (120lJ000 km^2^) of recently urbanized land in the U.S. subsuming natural forests. We leveraged a unique, well-characterized urban-to-rural and forest edge-to-interior gradient to identify the combined impact of these two land use changes - urbanization and forest fragmentation - on soil microbial community in native, remnant forests. We found evidence of mutualism breakdown between trees and their fungal root mutualists (ectomycorrhizal (ECM) fungi) with urbanization, where ECM fungi colonized fewer tree roots and had less connectivity in soil microbiome networks in urban forests compared to rural forests. However, urbanization did not reduce the relative abundance of ECM fungi in forest soils; instead, forest fragmentation alone led to strong reductions in ECM fungal abundance. At forest edges, ECM fungi were replaced by plant and animal pathogens, as well as copiotrophic, xenobiotics-degrading, and nitrogen-cycling bacteria, including nitrifiers and denitrifiers. Urbanization and fragmentation interacted to generate “suites” of microbes, with urban interior forests harboring highly homogenized microbiomes, while edge forests microbiomes were more heterogeneous and less stable, showing increased vulnerability to low soil moisture. When scaled to the regional level, we found that forest soils are projected to harbor high abundances of fungal pathogens and denitrifying bacteria, even in rural areas, due to extreme, widespread forest fragmentation. Our results highlight the potential for soil microbiome dysfunction - including increased greenhouse gas production - in temperate forest regions that are subsumed by urban expansion, both now and in the future.

**Significance Statement:** Urbanization and forest fragmentation are increasingly altering Earth’s ecosystems, yet the effects on soil microbiomes, crucial for plant health and climate regulation, remain unclear. Our data indicate that, in forested land, these two combined, compounding stressors reshape the soil microbiome in ways that could lead to more pathogen infections of plants and animals, higher rates of N loss due to denitrification, and the possibility of tree symbiont extinctions. By identifying the specific environmental stressors that lead to these microbiome shifts, our analysis can be used to inform urban development and forest management plans to mitigate impacts on the soil microbiome to sustain environmental quality and the ecosystem services that remnant native forests provide to society in the coming decades.

**Classification:** Biological Sciences/Ecology

## INTRODUCTION

The world’s forests are being rapidly urbanized and fragmented by land development^1,2^, with more than 70% located within 1 km of a forest edge^3^, especially in temperate regions which contain 52% more edge forest area than tropical regions^4^, but we are still in the early stages of understanding the impacts of urbanization and fragmentation on the important forest ecological communities. Soil microbes are critical regulators of carbon (C) loss and sequestration in forests, storing C belowground as stable microbial products^5^ and sustaining aboveground forest productivity by increasing nutrient availability for plants and suppressing pathogen loads^6^. Recent studies show dramatic shifts in the composition and activity of soil microbiomes with urbanization^7–10^ and forest fragmentation^11^, but the combined, compounding impacts of these two intense land development activities on soil microbial communities are unclear. Urbanization and fragmentation are expanding worldwide, with the loss of interior forests occurring up to seven times faster than the loss of edge forests in temperate deciduous forests^12^. At the same time, society has increased reliance on temperate forests in urban and rural areas for recreational, aesthetic, and health benefits, especially within the last few years^13,14^. Understanding the form and function of microbial communities within fragmented urban forests will become increasingly critical as we depend on them more heavily to sustain environmental quality and human well-being in the coming decades.

The multiple, interacting environmental stressors of urbanization and fragmentation may be particularly detrimental to microbial mutualists of plants, similar to the negative impact of environmental stress on plant-animal mutualisms^15^ and microbial symbionts of animals^16^. Urban areas have higher ambient air temperatures, greenhouse gas concentrations (e.g., carbon dioxide; CO_2_), nitrogen (N) loads (e.g., ammonium and nitrate deposition), and ambient pollutants (e.g. particulate matter and ozone; O_3_) relative to rural areas ^17–21^. Especially sensitive to these conditions are fungal symbionts of tree roots (e.g., ectomycorrhizal (ECM) fungi), which provide nutrients (such as N) to host trees in return for plant C^6^. ECM fungi often decline when soil N availability increases or plant photosynthesis slows due to land cover change or pollutant exposure, potentially because plants allocate less C belowground ^22–24^. ECM colonization of tree roots is lower in city streets compared to rural forests^25^, suggesting a potential tree-ECM mutualism breakdown under the stressful urban environment. A decline in ECM fungi in urban forests could open up trees to colonization by pathogens, as ECM fungi can provide pathogen protection to trees, and recent observations also indicate that foliar pathogen loads can be higher in urban compared to rural forests^26^. Fragmentation can intensify urban stressors by increasing the extent of forest edge^3^, which has higher air and soil temperatures^27^ and higher depositions of nutrients and pollutants compared to forest interiors^28^. If ECM fungi decline at forest edges, it could provide a greater opportunity for pathogenic taxa to dominate the soil community and diminish plant N uptake in fragmented, urban forests. However, some studies report increased plant C allocation to mycorrhizal fungi under elevated temperatures and drought conditions^29^, casting uncertainty on how plant mutualists respond to the combined influences of urbanization and edge effects.

Another way in which forest fragmentation may impact the soil microbiome is by increasing the vulnerability of soil communities to both abrupt and long-term environmental stress, similar to edge effects on forest trees. In temperate deciduous forests, trees grow faster and increase N uptake, which supports higher aboveground biomass at forest edges compared to the interior^4,30,31^, yet trees at forest edges are also more sensitive to heat stress^30^. Tree growth declines more rapidly after heat waves in urban forests than in rural forests, raising concerns that climate warming will have a disproportionately negative impact on C sequestration in highly fragmented urban forests^30^. Evidence is accumulating that there is also a growth-stress tolerance trade-off in microorganisms^32–35^, similar to that in macrorganisms^36,37^, such that a fast-growing soil community may also have a lower tolerance to environmental stress. When elevated N levels are applied experimentally to soils, they tend to initially select for fast-growing, copiotrophic microbes^38,39^, while often lowering total microbial diversity^40^. If fragmentation selects for fast-growing microbes at forest edges, these communities may have a lower tolerance to environmental stress, resulting in greater turnover - or instability - over time^41^. The effect would be exacerbated if soil microbiomes become more homogeneous with urbanization, similar to how urbanization can lead to more homogeneous plant and animal communities^42,43^. Urbanization can create unique environments, such as wet soils in dryland areas, which favor the growth of certain species^42–44^, and some have suggested the existence of an “urban suite” of soil microbes^45^, similar to groups of plants that often characterize urban green spaces^46^. However, in some instances, urbanization can generate high spatial habitat and community heterogeneity^47,48^, further accelerating the species evolution^49^, so it is unclear what the overall impact of urbanization and fragmentation have on microbial community structure if urban forest communities can withstand additional environmental stressors from edge effects, and what the role of these communities is in the health and functioning of forests.

There is reason to believe that soil microbes living at urban forest edges may have less C and nutrient sequestration capacity than their rural, interior forest counterparts. ECM fungi dominate soils in northern forests and can act as keystone taxa, providing C and N to the rest of the soil microbiome^50^, and controlling forest productivity to a degree that impacts atmospheric CO_2_ levels and the stability of the global climate^51^. The presence of ECM fungi is also potentially critical to increasing soil C stocks, because of the decomposition-resistant C in their biomass ^52–54^ and their ability to suppress opportunistic soil microbial taxa ^55–57^, such that a loss of ECM fungi could account for low soil C stocks at forest edges^58^. Urbanization may compound the impact of fragmentation on soil C sequestration by suppressing microbial activity: soil respiration rates are >40% lower at the edges of urban forests compared to rural forests^59^. However, high nitrification rates at forest edges^31^ indicate the potential for denitrifying microorganisms to thrive, which could lead to a switch from C- to N-based greenhouse gas production (e.g. N_2_O, NO_2_) at urban forest edges. Current ecosystem models do not include nitrogenous gas production from urban forest soils, and incorrectly predict increased soil CO_2_ release with urbanization^60^, such that resolving the uncertainty around soil microbial responses to urbanization and fragmentation may improve both our conceptual and predictive biogeochemistry models for large swaths of terrestrial forests.

Here, we examine the combined impacts of urbanization and forest fragmentation on soil microbial community composition and functional potential using the Urban New England (UNE) study^31,59^. UNE includes a series of eight fragmented, ECM-dominated temperate forest sites along a 120 km urban-to-rural gradient in the State of Massachusetts (MA), one of the most densely populated and densely forested states in the U.S. In MA, forests cover 60% of the land surface but are heavily fragmented due to urban sprawl^61^, where over two-thirds of the population lives^62^. Increases in temperature at the forest edge are comparable to the amount of warming this region experienced between 1900 and 1999 in response to climate change^63^. At UNE, duplicate sampling points are located along a 90 m transect from the forest edge to the forest interior at each site (Fig S1, Table S1), a design that allows us to uniquely capture the interacting effects of forest fragmentation and urbanization on a widespread and ecologically important forest type within the Northern Hemisphere.

We hypothesized that urbanization and forest fragmentation interact to reduce ECM abundance in soils, shifting soil microbial communities towards more pathogenic and copiotrophic taxa, particularly at urban forest edges. We also expected urbanization and fragmentation to homogenize soil microbial communities, selecting for communities with fewer associations between taxa and making soil communities more vulnerable to environmental stress, which could explain changes in soil N cycling and C storage observed at UNE^31,58^. To test these hypotheses, we paired three years of data (across 2018, 2019, and 2021) on soil microbial community composition with soil biogeochemistry (e.g. full elemental analysis, soil respiration rates, and rates of N cycling) and physiochemistry (e.g., pH, temperature, moisture) data from the same soil samples, as well as data on root biomass, ECM colonization rates, plant density, and community composition. We tested the impact of urbanization (measured as the distance from an urban center (Boston Common in Boston, MA) and fragmentation (classified as the distance from the forest edge^31,59^) on soil microbial communities using a combination of linear regression models that accounted for spatial autocorrelation in community composition (Fig. S2), as well as differential abundance analyses and network construction. To understand the potential roles of microbial groups on soil C and N cycling, we also looked for relationships between the genomic capability of the microbial community to process soil C and N and total soil respiration rates^59^, N transformations^31^, and C stocks^58^. Additionally, we scale the results of our analyses to the state level, leveraging soil microbiome data we generated for each of the ECM-dominated USDA FIA forest types in Massachusetts.

## RESULTS AND DISCUSSION

In contrast to our expectations, neither urbanization nor its interaction with fragmentation reduced ECM fungal abundance in soils of remnant, native forests. Instead, fragmentation alone had striking negative effects on ECM fungi: ECM fungi declined precipitously in soil at the edges of both urban and rural forests (Fig 1a) despite a similar % ECM trees and similar root biomass at forest edges compared to forest interiors (Fig. S3). This result contrasts with recent reports of reduced ECM fungal abundance in urban soils worldwide^9^, which could be explained in part by edge effects on fragmented urban landscapes. Loss of ECM fungi was correlated with an increase in fungal and bacterial diversity in soils at forest edges (Fig. S4) due to an increase in pathogenic microbes (Fig. 1b, c, Table S2), xenobiotics-degrading and copiotrophic bacterial communities, as well as nitrate-producing and consuming bacteria (Table S2, Fig. S4). This shift in soil microbiome composition from ECM fungi to nitrate-cycling bacteria was mirrored by increased nitrification rates at forest edges (Fig. S5)^31^ and parallels results from other studies showing that nitrifying bacteria may compete with ECM fungi for ammonium^57^. Denitrifying bacteria - including those that conduct partial denitrification - also show high abundance at the forest edge, increasing the potential for forest fragmentation to release nitrogenous greenhouse gas (e.g., N_2_O and NO_2_) that may accelerate atmospheric warming and feedback to increase drought stress in forest edges. Interestingly, animal pathogen abundance is also high at forest edges (Fig 1b, Table S2), as is the abundance of dung-saprotrophic fungi (Table S2), suggesting the greater influence of animals on the soil microbiome at both urban and rural forest edges.

**Fig. 1.**
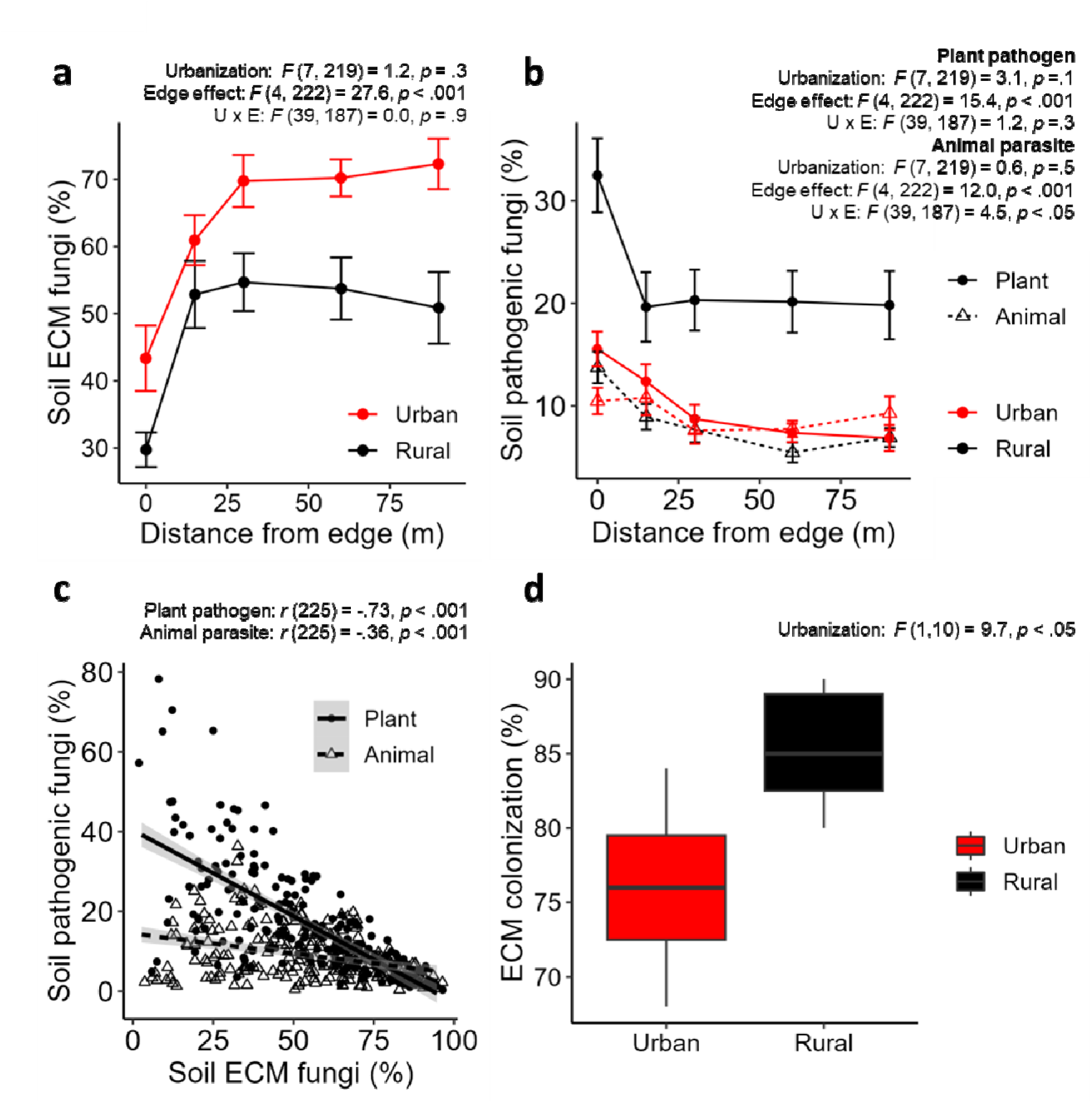
Relationships between urbanization and fragmentation and key plant- and animal-associated soil microbes. The relationship between distance from the forest edge and the relative abundances of (a) ECM fungi and (b) pathogenic fungi are shown. (c) ECM fungi are anti-correlated with pathogenic fungi in the soil, including animal pathogens. Differences in (d) ECM colonization of oak trees between urban sites and rural sites illustrate the breakdown of tree-ECM mutualisms with urbanization. ANOVA results for the linear mixed-effect models testing for relationships with urbanization (negative distance from Boston), edge effects (negative distance from edge), and their interactions (U×E) are shown for ECM fungi (a), fungal pathogens (b), and ECM colonization (d). Correlation test results for the fungal group are shown (c). The error bar shows the standard error. Asterisks represent *P* values (\**P* < 0.05, \*\**P* < 0.01, \*\*\**P* < 0.001).

Despite high relative abundances of ECM fungi in urban forest soils, we found evidence of tree-ECM mutualism breakdown under urbanization: urban trees had lower ECM colonization rates on roots (Fig.1d), despite higher ECM tree and root abundance (Fig. S3) and increased plant N uptake (leaf N concentration, Fig. S5, Table S2) relative to rural trees. This result is consistent with the one other study that quantified ECM colonization of trees planted in urban street tree pits vs. rural forests^25^, which found consistently lower ECM colonization of trees in street pits. High rates of net N mineralization in urban sites^31^ might explain this phenomenon, as we found that experimental N addition at rural forests reduced the ECM colonization rate to levels close to those found in urban forests (Fig. S6). High levels of inorganic N in soil have an overwhelmingly negative effect on tree-ECM symbioses^64^, suggesting that the benefit of strong tree-ECM mutualisms under the high temperature and drought stress in urban systems might not outweigh the benefit of reducing C allocation to symbionts under high soil N availability.

Tree-ECM mutualism breakdown, combined with additional unique environmental conditions in urban forests, changed community structure and species connectivity in the soil microbiome. In contrast to our expectations, network analysis revealed that urban soil microbiomes were highly connected, with taxa clustering into densely connected ‘cliques’ (Fig. 2a, b) that indicate strong covariance of taxa in response to environmental conditions^65^. Interestingly, ECM fungi were less central in urban soil community networks than in rural networks (Fig 2d), indicating lower activity of this group in urban environments. Instead, the most connected taxa in urban soil networks were other, less well-studied bacteria and fungi that have a large genomic capacity for diverse C and N metabolism (Figs 2, S7). This switch from soil microbiomes connected by tree symbionts to ones potentially structured by the activity of free-living saprotrophs suggests that the combination of higher soil temperatures and release from ECM activity may be the major impact of urbanization on microbiome structure and metabolism. Microbial network connectivity and complexity were positively correlated with soil and foliar C content (Fig. S5), suggesting a strong environmental selection of microbes by these resources, which has also been found in other studies^66,67^. Complex networks of microorganisms in urban forest soils may also feedback to contribute to the ecosystem functions observed there^68^, such as unusually high ammonification rates (Fig. S5).

**Fig. 2.**
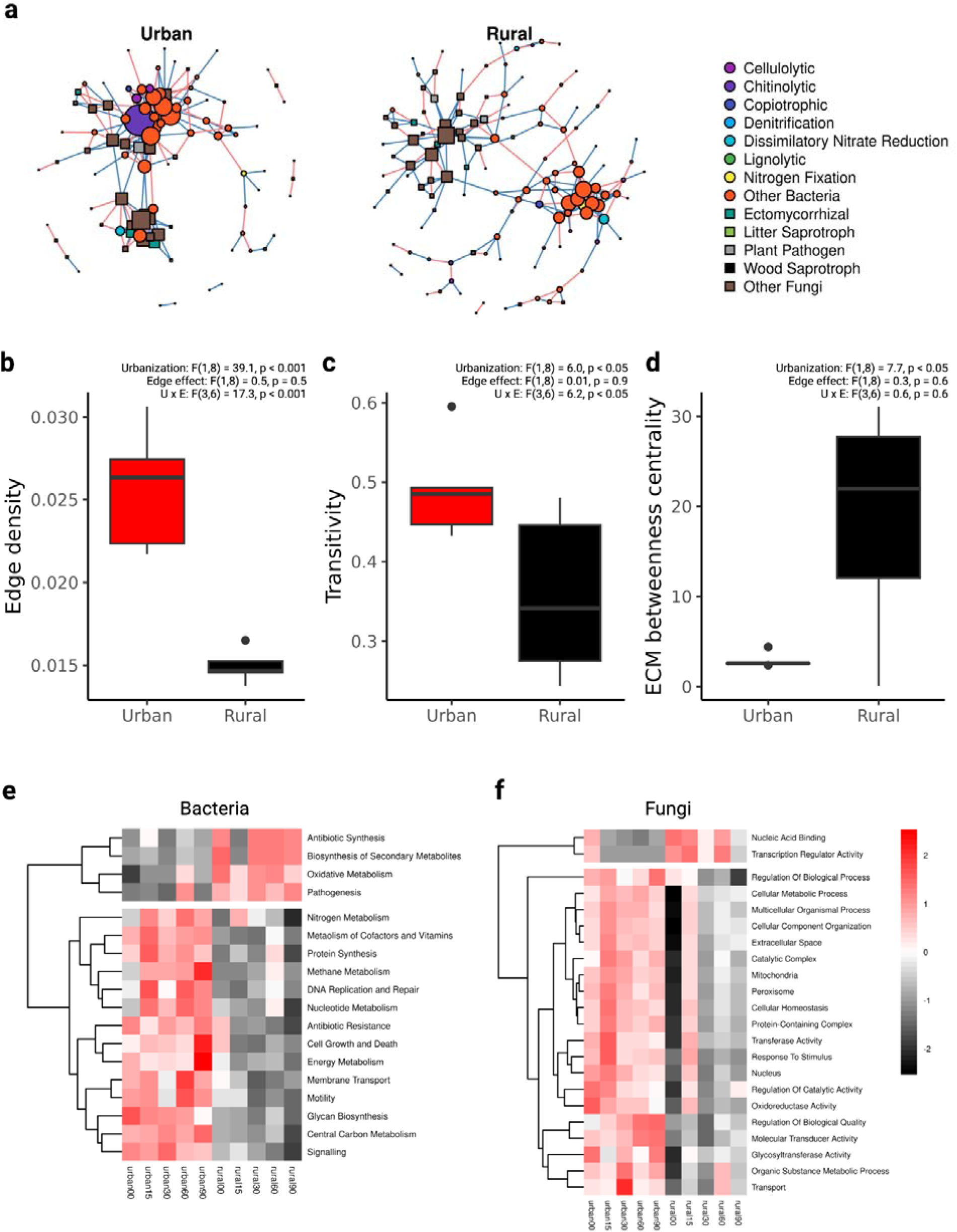
(a) Networks of urban and rural forests. Circles represent bacterial functional groups and squares represent fungal functional groups. Sizes of circles and squares represent the degree, or number of connections, of the taxa. Positive associations are colored blue and negative associations are colored red. ANOVA results for the linear mixed-effect models testing for relationships with urbanization (negative distance from Boston), edge effects (negative distance from edge), and their interactions (U×E) are shown for network metrics edge density (b), transitivity, or ‘cliqueness’ (c), and ECM betweenness centrality, or network importance (d). Heatmaps show the bacterial (e) and fungal (f) genetic pathways enriched in urban and rural networks. Pathway abundances are scaled to z-scores by row in the heatmap. Pathways above the heatmap breaks in both (e) and (f) are enriched in rural networks, while pathways below the heatmap breaks are enriched in urban networks. Bacterial gene groups in enriched pathways can be found in Fig. S7.

Together, urbanization and fragmentation interacted to generate “suites” of soil microbes that characterized each forest type that was phylogenetically diverse (Fig. S8) but functionally constrained. For example, urban forest interiors were characterized by ECM fungi and cellulolytic, nitrate/nitrite-reducing, and methanotroph bacteria, while urban edges were defined by plant pathogens and nitrifying taxa (Fig. 3, Fig. S9). Consistent with our expectations, urban forest soil communities were more homogeneous than rural forests, but only in forest interiors (Fig. 4a). This phenomenon corresponded to the homogenization of soil properties (soil temperature, moisture, pH, organic matter content, ammonium content, and nitrate content) in the interiors of urban forests (Fig. 4b), similar to other studies that have found urbanization homogenizes the physical environment^69^ and contributes to biotic homogenization of plants and animals^42,44^. By contrast, forest edge microbiomes were more heterogeneous and tended to change more dramatically - or be less stable - over time than at the forest interior (Fig 5a). Instability of both bacterial and fungal communities was linked to increased vulnerability to low soil moisture (Fig. 5b), which is most severe at the forest edge^58^. Soil microbial vulnerability to drought has been hypothesized for the last decade^70^ and our data provide the first evidence that this is a key way in which urbanization and fragmentation impact the soil microbiome. Interestingly, bacteria are less stable than fungi and more sensitive to low soil moisture at forest edges (Fig 5b), potentially because they are more dependent on water for dispersal^71,72^. Forest edges also experience larger fluctuations of soil water content than interiors, because of high light intensity in edge habitats^73^, a process that is known to change microbial community composition^74,75^. In addition, because ECM fungi can connect microbiome members by providing plant C and N^50^, a loss of ECM fungi could also partially explain community instability at forest edges (Fig. 5a).

**Fig. 3.**
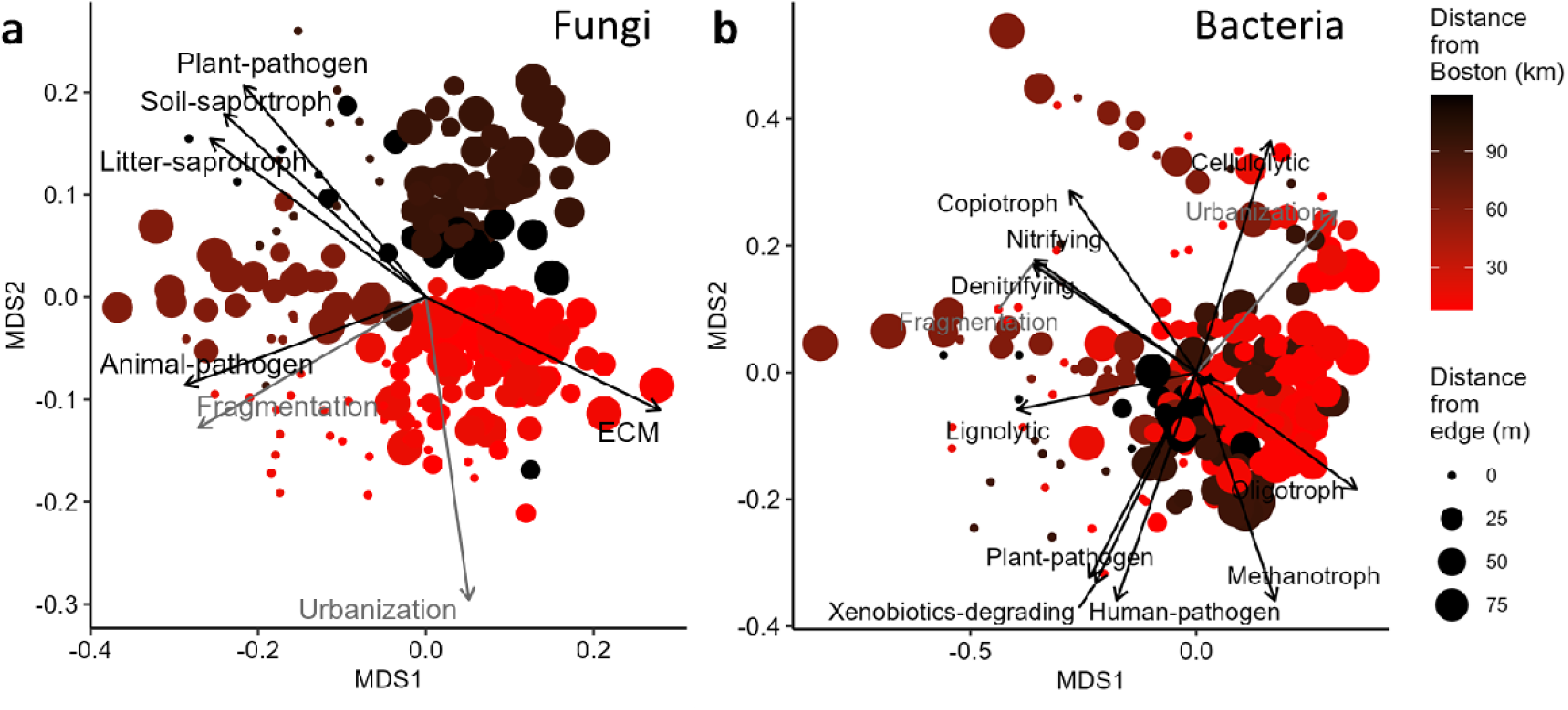
Community composition of (a) fungi and (b) bacteria, including “suites” of urban/rural and edge/interior forest soil microbes. Point color and size represent the distance from Boston (km) and the distance from the forest edge (m), respectively. Differential abundance of microbial groups across suites was tested using the vegan::envfit function^105^, and significant groups are shown on each plot^105^.

**Fig. 4.**
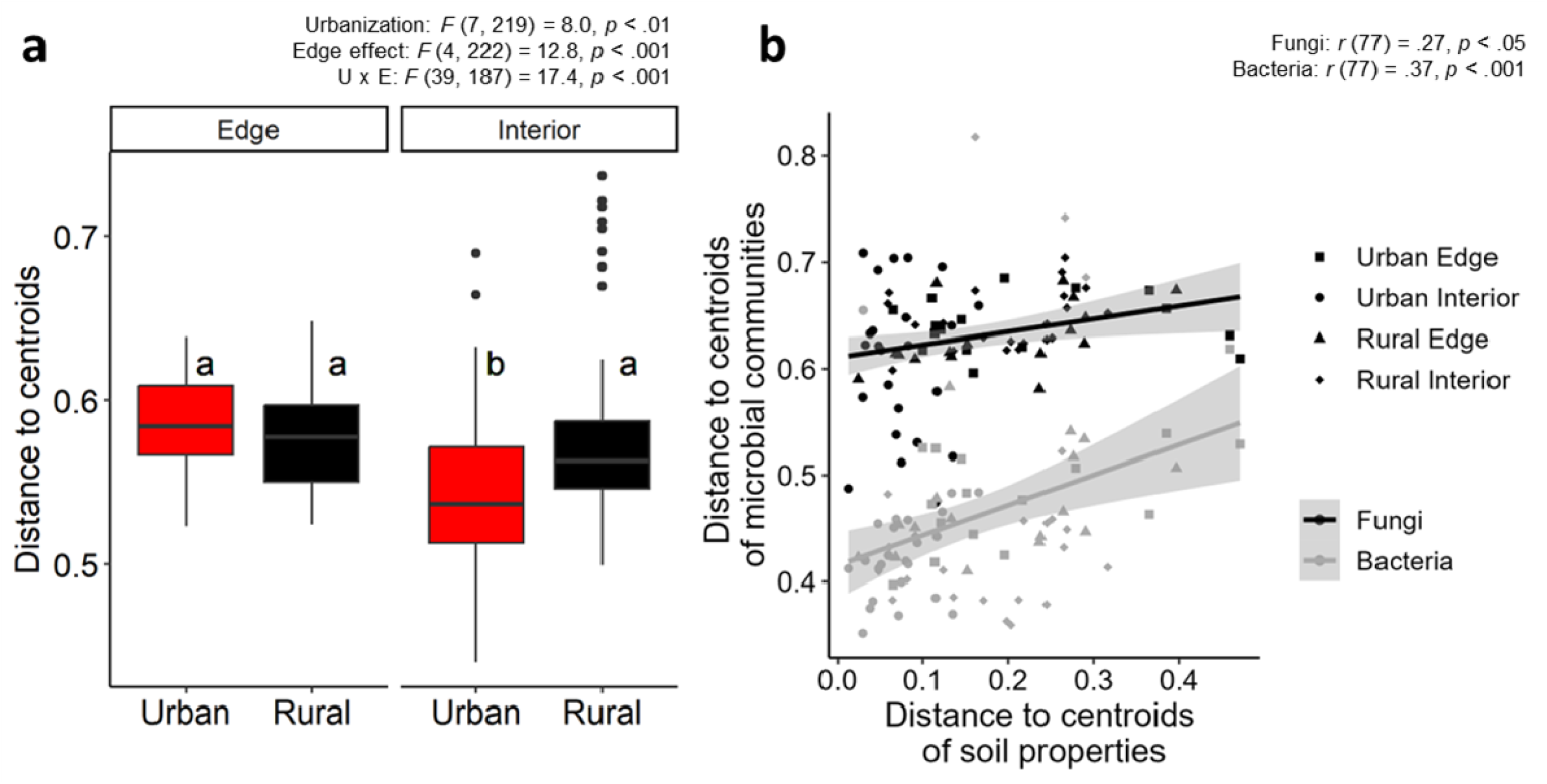
(a) Homogenization (measured as distance to centroids) of soil microbial (both fungal and bacterial) communities in each location (urban edge, urban interior, rural edge, rural interior), and (b) the relationships between homogenization of soil properties and soil microbial communities. The result of ANOVA for the linear mixed-effect model for urbanization, edge effects, and their interactions (U×E) and the result of Tukey’s multiple comparison tests (a), and the correlation coefficients (b) are shown. Asterisks represent *P* values (\**P* < 0.05, \*\**P* < 0.01, \*\*\**P* < 0.001).

**Fig. 5.**
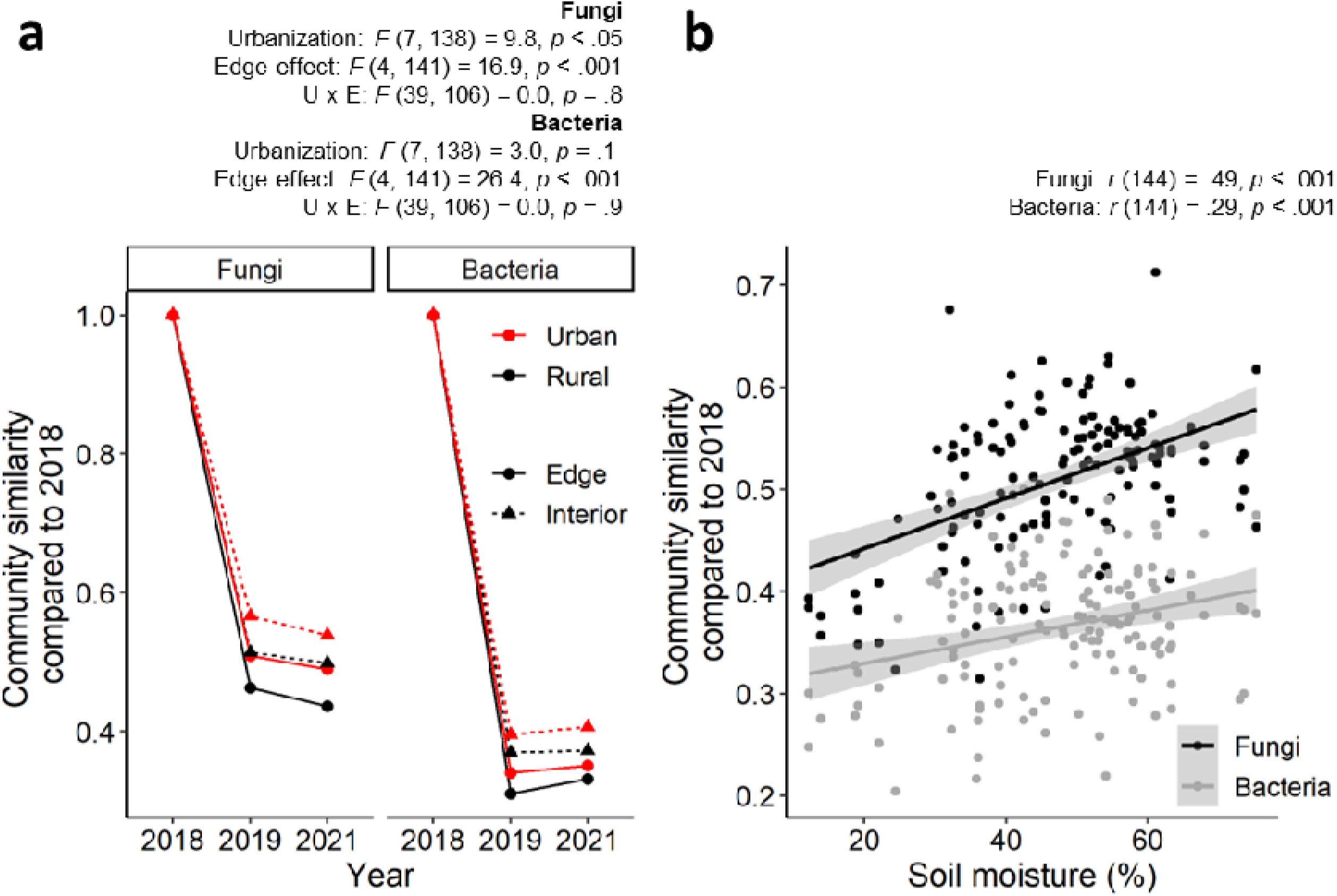
Community instability across urban/rural and edge/interior forest soils (a) and community vulnerability to drought stress across sites (b). Instability was calculated as community similarity (calculated as Aitchison distance) compared to the 1st year (2018), of which community was the baseline of the comparison (a), and vulnerability to drought stress was calculated as the relationship between community instability and soil moisture (b). The result of ANOVA for the linear mixed-effect model for urbanization, edge effects, and their interactions (U×E) is shown (a), and the correlation coefficients for fungal and bacterial community instability are shown (b). Asterisks represent *P* values (\**P* < 0.05, \*\**P* < 0.01, \*\*\**P* < 0.001).

When we scale the change in forest soil microbiomes with urbanization and fragmentation across ECM-dominated forests for the state of Massachusetts, we project unexpectedly high regional loads of plant and animal pathogens (Fig. 6). While ECM fungi comprise > 55% of the soil fungal community for forests in most of this territory, plant pathogens make up over a quarter of the soil fungal community in over 77,000 ha of forest area. This phenomenon is associated with widespread forest fragmentation, resulting in strikingly high projected levels of fungal pathogens in the rural, typically regarded as pristine, western part of the state. The loss of tree symbionts and the rise of pathogenic microbiomes in forests could explain reduced tree growth at the edges of temperate forests that experience extreme heat stress^30^. In addition to fungal pathogens, we project extremely high levels of denitrifying bacteria across the state: over 130,000 Ha of forest across the state harbors more than double the relative abundance of denitrifiers found in soils of intact, interior rural forests (Fig. 6), indicating potential for greater N-based greenhouse gas emissions from soil than previously expected.

**Fig. 6.**
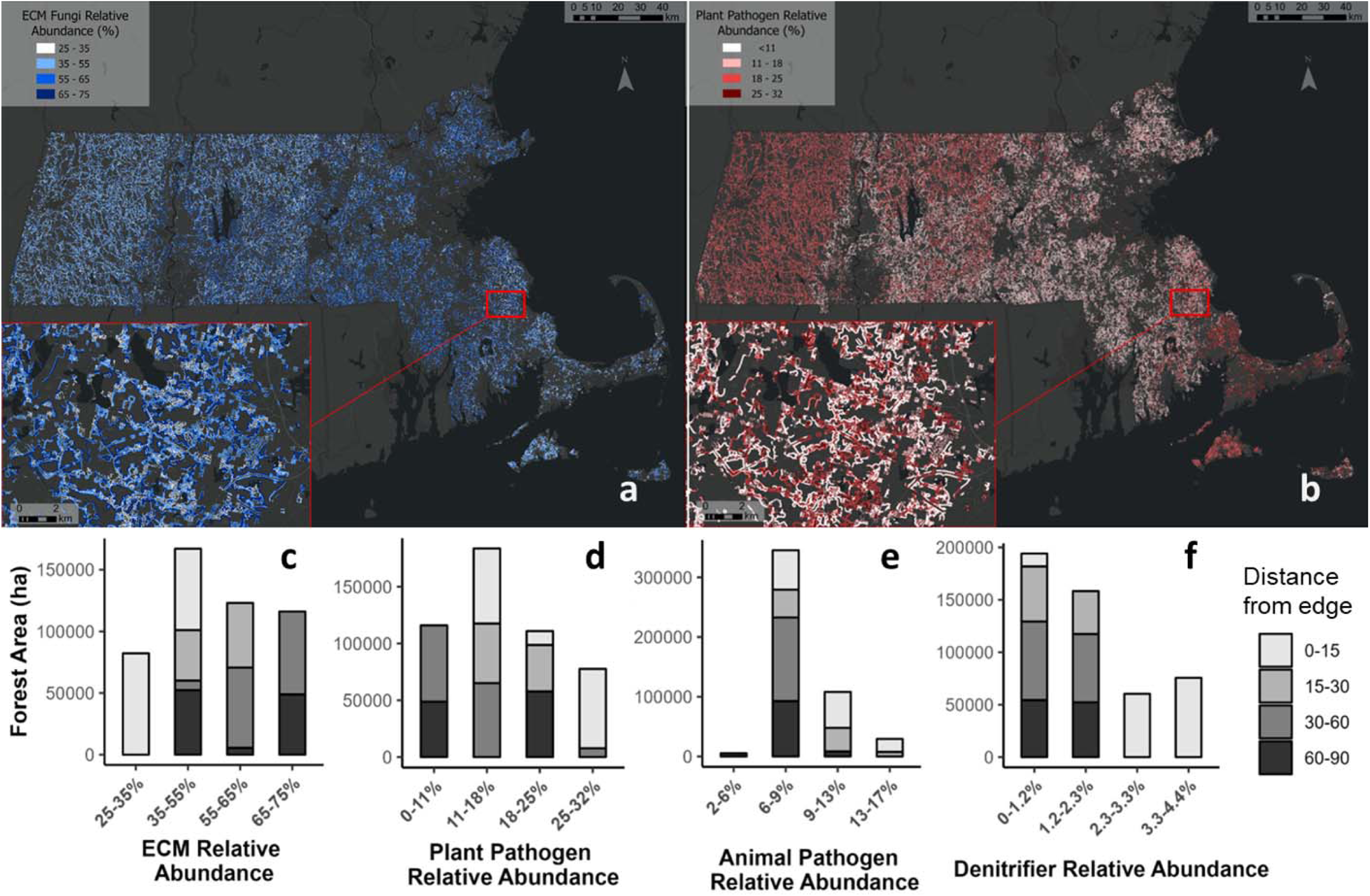
Projected abundances of microbial groups in ECM-dominated forests in mixed rural and urban landscape across the state of Massachusetts. Forest type groups at UNE field sites were matched with US Forest Service forest types, then relative abundances of key fungal (ectomycorrhizal fungi (ECM, a, c), plant pathogens (b, d), and animal pathogens(e)) and bacterial denitrifying groups (f) calculated for each UNE plot were projected across interior and edge forest area within the state. Edge forest was categorized as forest area bordered by non-forest in distance-from-edge bins (15 m, 30 m, 60 m, 90 m).

## CONCLUSION

Arguably one of the greatest – and most poorly understood - threat to temperate forests is fragmentation through land clearing and development. Northern temperate hardwood forests are already the most urbanized and fragmented forest type in the world and have the highest likelihood of becoming more fragmented and urbanized in the coming decades^1,2^. Our study suggests that urbanization and edge effects from this land development have strong negative impacts on tree symbionts and their role as keystone taxa in soil microbial communities. In contrast to our expectations, urban forest soils are dominated by ECM fungi, because of the unexpectedly high percentage of ECM trees in our urban sites. However, higher soil N availability - driven by increasingly warmer soil temperatures and high N pollution in urban areas - may further threaten tree-ECM symbioses and dramatically change urban forest composition in the near future^76^. Our observation that soil at forest edges has fewer ECM fungi and increases in pathogenic and stress-tolerant microbes indicate that where ECM trees decline, we may see additional increases in pathogen and denitrifier loads beyond what we now project for the region.

## MATERIALS AND METHODS

### Study system and sample collection

The Urban New England (UNE) study consists of eight mid-successional, temperate, deciduous forest sites spanning an urban-to-rural gradient in Massachusetts (Fig. S1, Table S1). Here, we defined the urbanization gradient as the distance from an urban core^31^, where Park Street Station in Boston represented the urban end of the gradient, and Leverett, MA represents the rural end. Microbial communities are more similar when grouped based on the distance from the urban center (Boston), rather than by other well-known urbanization indicators, such as impervious surface area (Figs. 3, S10)^77^. Each site at UNE consists of 5 plots (40 plots in total per site) that are 10 m × 20 m in area and are located at 0, 15, 30, 60, and 90 m from the forest edge. The climate at these sites is humid continental with warm summers (mean monthly temperatures of 18.6°C to 21.7°C) and cold, snowy winters (−4.3°C to −0.1°C), and 1100-1300 mm of precipitation distributed evenly throughout the year (NOAA 2021). Dominant tree species are mostly ECM-associates such as red oak (*Quercus rubra*), white pine (*Pinus strobus*), sweet birch (*Betula lenta*), although some plots have a high abundance of red maple (*Acer rubrum*), and sugar maple (*Acer saccharum*). The total tree basal area per plot ranged from 10.5 to 94.0 m^2^ ha^-1^ (Table S1). Additional site details are available in Garvey et al (2022)^59^ and Caron et al. (2023)^31^.

Organic horizon soils (0.8 – 9.7 cm depth) at UNE were collected in August 2018, July 2019, and July 2021. Two replicate soil samples were collected in each plot at the same time and location as soil respiration and the temperature measurements were taken from PVC respiration collars installed on the forest floor^59^ (227 samples in total; 40 plots × 2 replicates × 3 sampling years). We omitted 1 and 12 samples from 2018 and 2021, respectively, because of plot accessibility. Soil samples were kept on ice during same-day transport back to the laboratory, stored at 4 lJ until processing, and processed within 72 hours of sampling. Soils were sieved through a 2mm sieve and then a subsample was stored in a −80 lJ freezer until DNA extraction. Additional fresh soil subsamples were used to measure soil physicochemical properties such as moisture, pH, nutrient content, and fluxes^31,59^. Analysis details for all biogeochemical and plant measurements can be found in the Supplementary Methods.

### Soil DNA extraction, quantitative PCR, fungal/bacterial amplicon sequencing, and bioinformatics

<u>DNA extraction</u>: Total soil DNA was extracted from approximately 0.25 g of soil using the DNeasy PowerSoil HTP 96 Kit (QIAGEN, Hilden, Germany). Two technical replicates were extracted from one soil sample, resulting in 454 extracts in total ({8 sites × 5 plots × 2 biological replicates × 3 sampling years −13 samples} × 2 technical replicates). Extracts were stored frozen at −80lJC until further analysis.

<u>Quantitative PCR (qPCR):</u> The two DNA extractions (technical replicates) from each soil sample were pooled to better represent soil microbiome diversity^78^. Modified versions of the primer set fITS7 and ITS4 were used for fungi, amplifying the ITS2 region of rDNA^79,80^ and modified versions of the primer set 806R and 515f, which target the v4 16S region of rDNA, were used for bacteria^81^. Additionally, an abundance of AM fungal (18S rDNA), N-fixing bacterial (nifH), nitrifying bacterial (amoA), and denitrifying bacterial (nosZ) genes were quantified^82–85^. Detailed descriptions of qPCR conditions are outlined in Supplementary Methods.

<u>Amplicon sequencing:</u> the same fungal and bacterial primer sets used for qPCR were used for amplicon sequencing, with the exception that primers contained both the Illumina adapter and individual sample indexes^86,87^. Amplicon sequencing was conducted on the pooled DNA extract per sample (227 samples total). Amplicons were checked by agarose gel electrophoresis, then cleaned using Just-a-Plate 96 PCR Purification and Normalization Kit (Charm Biotech, MO, USA) and quantified using Qubit HS-dsDNA kit (Invitrogen, Carlsbad, California, USA). 16S and ITS amplicons for each sample were mixed at equimolar concentrations. Both 16S and ITS amplicons for 80 samples at maximum were combined into a single library for sequencing, generating six libraries in total. Each library was subject to 250 base pair (bp) paired-end sequencing on Illumina MiSeq run at the TUFTS Genome Sequencing Core facility. Sequence data were deposited in the Sequence Read Archive at NCBI under accession number DRA015736.

Bioinformatics was performed in the R software environment (version 4.0.0), where the R package dada2 was used for sequence quality control, paired-end assembly, identification of amplicon sequence variants (ASVs), and taxonomy assignment^88^. Taxonomy was assigned to ASVs using the naive Bayesian classifier method^89^ in combination with the UNITE database (v. 7.2)^90^ as the reference for fungal ITS ASVs and the SILVA database (release 138.1)^91^ as the reference for bacterial ASVs. To assign taxa to functional guilds, fungal ASVs were searched against the FungalTrait database^92^ and bacterial genera were searched against an in-house functional database created based on substrate enrichment experiments and the presence of genes coding for enzymes in specific biochemical pathways (copiotroph, oligotroph, cellulolytic, ligninolytic, methanotroph, chitinolytic, assimilatory nitrite-reducing, dissimilatory-nitrite reducing, assimilatory nitrate-reducing, dissimilatory nitrate-reducing)^93,94^. “Suites” of bacteria and fungi were calculated for location group (i.e., Urban edge, Urban interior, Rural edge, and Rural interior) using indicator species analysis^95^. PICRUSt2^96^ was used for predicting the bacterial gene abundances for each sample based on KEGG Orthology (KO)^97^ annotations.

For network construction, we collapsed fungal and bacterial amplicon sequence counts to the genus level and normalized each genus to qPCR data for fungi (ITS) and bacteria (16S). Samples from the same plot for urban and rural forests (i.e., urban forest 0m plots, urban forest 15m plots, rural forest 0m plots, etc.) across all sampling years were combined to construct a network (2 samples per plot × 4 forests per forest type × 3 sampling years = 24 samples per network). Abundance tables were filtered (see Supplementary Methods) and networks were constructed using the SpiecEasi package of R^98^ (parameters are described in the Supplementary Methods). Networks were analyzed for measures of connectivity, overall topological structure, and taxon importance using the igraph package of R^99^. We also conducted a synthetic metagenome analysis of network taxa, using the KEGG Orthology (KO)^97^ annotations of genomes for bacterial taxa and Gene Ontology (GO)^100^ annotations for fungal taxa in each network, following a recent protocol for synthesizing fungal genomes from the Joint Genome Institute 1000 Fungal Genomes project^101^.

### Scaling to the State of Massachusetts

To explore the implications of our findings over a mixed rural and urban landscape, we scaled our results across the state of Massachusetts. We first applied forest compositional groups defined by the US Forest Service (Forest Inventory & Analysis User’s Manual Appendix D) to categorize each of our study sites. We used a one-meter resolution land-cover map developed by the Massachusetts Bureau of Geographic Information (available at https://www.mass.gov/info-details/massgis-data-2016-land-coverland-use) to categorize forest area bordered by nonforest into distance-from-edge bins (15 m, 30 m, 60 m, 90 m; *sensus* Reinmann et al., 2020^30^). We then combined the binned map of edge-influenced forest area with a 250 m resolution forest composition map^102^ to identify edge forests with species assemblages matching those of our study sites. Finally, we estimated the relative abundances of microbial community constituents in Massachusetts based on patterns of forest fragmentation and species composition.

### Statistical analysis

Prior to statistical analyses, data were normalized in multiple ways, depending on the data type. For qPCR, data were normalized via logarithmic function. For amplicon sequence data, we normalized ASV counts across samples using the rarefaction, which lowers the false discovery rate of ASVs when samples vary >10-fold in sequencing depth^103^. ASV counts per sample were rarefied by equalizing final read numbers to 18,028 reads for fungal communities and 11,384 reads for bacterial communities using random pick-up based on the minimum read number. We found that normalization by ANCOM-BC^104^ and rarefaction produced similar results across analyses (Table S3), so we report results of taxonomic and functional group analyses with rarefied data, allowing us to include rare taxa in the analysis.

To explore relationships between urbanization, edge effects, and the soil microbiome, we used distance from Boston and distance from the edge, as well as their interaction, as the fixed independent variables for all statistical models. To account for repeated sampling at individual locations, the site was included as a random variable in each model^31^. Shannon’s alpha diversity for the whole fungal and bacterial community was calculated by the vegan package^105^ in R. For community instability, Aitchison’s dissimilarity was calculated^106^. To test for urbanization and edge effects on community homogenization, the distance to centroids was calculated in each location group (i.e., Urban edge, Urban interior, Rural edge, and Rural interior). Urban and rural site designations were based on distance from Boston^31^ and the edge included 0-15 m, while the interior included a 30-90 m distance from the forest edge. Urban and edge effects on microbial community were calculated using a linear mixed-effect model (LMM). ANOVA (analysis of variance) test for LMMs was run for values, standardized by the scale function in R, using the lme4 and lmerTest packages^107,108^ in R. Community differences among soil samples were visualized by nonmetric multidimensional scaling (NMDS) of community structure based on the Bray-Curtis dissimilarity index using the metaMDS function in the vegan package^105^ of R. The envfit function was used to illustrate significant correlations between fungal or bacterial functional groups and community dissimilarity among soil samples.

We measured the relationships between environmental factors, urbanization, and edge effects using an LMM. At UNE, urbanization was significantly positively correlated with soil temperature, and net N mineralization rate, but significantly negatively correlated with total tree basal area (Table S2b). Fragmentation was significantly positively correlated with soil pH, soil nitrate content, net nitrification rate, total tree basal area, and foliar C: N ratio, but significantly negatively correlated with soil moisture, SOM concentration, soil net N mineralization rate, and soil respiration rate (Table S2b). To see the significant correlations between microbial functional groups and environmental variables, *Pearson*’s correlation test was used. We set the level of significance at 5% for all statistical tests. The codes are available on GitHub (https://github.com/Chikae-Tatsumi/UNE).

## ACKNOWLEDGEMENTS

We thank the Arnold Arboretum of Harvard University, Harvard Forest, Massachusetts Department of Conservation & Recreation, Massachusetts Department of Environmental Protection, the Massachusetts Department of Transportation, Massachusetts Audubon Society, National Grid, Cities of Newton and Lexington, Massachusetts, and all landowners for allowing research on their properties. We are also grateful for the assistance in the laboratory, fields, or data analysis, or the helpful comments on earlier drafts of this manuscript from members of Bhatnagar, Templer, and Hutyra Labs, as well as Jerry Melillo, John Hobbie, Maggie Anderson, Talia Michaud, Takuro Ogura, Jonathan Gewirtzman, Stephen Caron, Nahuel Policelli, Corinne Vietorisz, Michael Silverstein, and Zoey Werbin. This study was financially supported in part by Grant-in-Aid for JSPS Research Fellow (Grant No. 17J07686, 20J00656), Oversea Challenge Program for Young Researcher (Grant No. 201980107) to C.T.; United States Department of Agriculture and National Institute of Food and Agriculture grant to Templer and Hutyra (USDA NIFA 67003-26615), DOE BER award DE-SC0020403, a Patricia McLellan Leavitt Research Award, a startup fund from Boston University and Boston University’s internal Peter Paul Professorship fund to J.M.B.; and the Boston University Microbiome Initiative Accelerator Program and NSF NRT DGE 1735087 to K.A. The genomics work conducted by the U.S. Department of Energy Joint Genome Institute, a DOE Office of Science User Facility, is supported by the Office of Science of the U.S. Department of Energy under Contract No. DE-AC02-05CH11231.

