## Supplementary material for "Urbanization and fragmentation interact to drive mutualism breakdown and the rise of unstable pathogenic communities in forest soil"

**Supplementary Methods**

**Soil biogeochemical and physicochemical measurements**

Physiochemical and biogeochemistry measurements were made on soil samples collected for DNA extraction and microbial community analysis (40 plots × 2 replicates × 3 sampling years). Gravimetric soil water content was measured by drying the soil at 65 °C for more than 48 hours. Soil pH was measured on a 1:2 soil/water suspension using a pH meter. Soil organic matter content was measured by combusting dried soil at 450 °C for 4 hours in a muffle furnace. Total soil C and N contents were analyzed on an NC2500 elemental analyzer (CE Elantech, Lakewood, NJ, USA). Soil soluble N was extracted with 2 mol/L KCl at a 1:10 fresh soil/extractant ratio. Soil net ammonification and net nitrification rates were measured *in situ* using the “buried bag” method^1–4^. More detailed information on N mineralization measurements is written in Caron et al (2023)^5^. Soil temperature was measured in field soil at each sampling time point using a Hanna Instruments Thermistor Thermometer (*Waterproof Thermistor Thermometer: HI93510N*, 2021) at 10 cm depth^6^. At the same time, soil respiration was also measured using an LI-COR LI-8100A soil respiration chamber system (±1.5% accuracy for CO_2_ reading; LI-8100A Specifications, 2021) from PVC soil respiration collars, which were 20.2 cm in diameter, mounted in the soil approximately 4 cm deep and extending aboveground roughly 4 cm^6^. We used the temperature and respiration values that were collected on the closest dates (< two weeks) to the soil collection dates. More detailed information is written in Garvey et al (2022)^6^ and Garvey et al. (in review)^7^.

**Leaf and root sample collection and measurement**

Samples for root colonization were collected at four of the UNE sites, including two urban sites (Hammond Woods, Sutherland Woods) and two rural sites (Harvard Forest 1 and Harvard Forest 2) in October 2018. Soil for root colonization was also collected at the Harvard Forest Chronic N Amendment Experiment, which ran from 1988 to 2019 and includes three 30 × 30m plots: control which received no N addition, low N addition (50 kg N ha^-1^ yr^-1^), and high N addition (150 kg N ha^-1^ yr^-1^) with the N addition treatments applied in the form of ammonium nitrate (NH_4_NO_3_)^8^. Samples were collected near three oak trees at the UNE plots (at 40, 60, and 80 m from the forest edge) and at the Harvard Forest Chronic N Amendment Experiment across each of the plots. At each oak tree that was sampled, three 10 × 10 cm soil blocks of the O layer were collected 0.5 m away from the tree in 3 equally spaced spots around the tree base. Samples were pooled, resulting in one sample per forest plot. For each tree, a composite subsample was soaked in water and then all roots were collected and cut into 2 cm lengths. Fifty 2 cm lengths of fine root were selected and analyzed under a dissection scope. Each root tip was identified as either not colonized or colonized by ectomycorrhizae (presence of at least one hyphal mantle = colonized). Data are reported as percent root colonized per soil sample.

For measuring the root density, 10 cm depth soil cores (2.4 cm radius) were collected with a hammer core from near each PVC collar at each plot in 7 of the 8 UNE sites (site HF05 was omitted due to accessibility issues) in August 2021 (n=147 in total). Soil core depth was measured in the field and used to calculate root density in units of g root per cm^3^ soil. Root biomass was measured by washing the separated roots in a 2 mm metal sieve with DI water. Each root was picked from the sieve and placed in its respective coin envelope. The roots were dried for 72 hours at 60°C and weighed.

Foliar samples were collected from oak trees using a pole pruner and approximately 3 trees were sampled at each plot, though plot limitations sometimes prevented a full collection of samples from each site. At the forest edge, sun leaves were collected from the edge side of the trees. In the forest interior, sun leaves were collected from gaps in the canopy that received sunlight. Leaves were then separated from branches at the base of the petiole and the entire sample (petiole included with the leaf) was ground, then transferred to a glass scintillation vial. Prior to weighing, sample vials were placed in the drying oven at 50℃ with caps loosened to allow moisture to vent. The C and N concentrations of foliar samples were measured on an NC2500 elemental analyzer (CE Elantech, Lakewood, NJ, USA). Additional information can be found in Caron et al. 2023^5^.

**Quantitative PCR**

The quantitative PCR of microbial groups followed published protocols^9^ with modifications. The PCR was carried out in 20 µl reactions, which included 2 µl of 20-fold diluted genomic DNA, 0.8 µl of each 10 µM primer, and 10 µl of PowerUp™ SYBR™ Green Master Mix (Thermofisher Scientific, Waltham, MA). When the 20-fold diluted DNA did not amplify, 100-fold diluted DNA was used. The primer sets used for the qPCR were as follows: 515f (GTG CCA GCM GCC GCG GTA A) - 806R (GGA CTA CHV GGG TWT CTA AT) for bacterial 16S rDNA^10^, fITS7 (GTG ART CAT CGA ATC TTT G) - ITS4 (TCC TCC GCT TAT TGA TAT GC) for fungal ITS rDNA^11,12^, AMG1F (ATA GGG ATA GTT GGG GGC AT) - AM1 (GTT TCC CGT AAG GCG CCG AA) for arbuscular mycorrhizal fungal 18S rDNA^13,14^, nifHF (AAA GGY GGW ATC GGY AAR TCC ACC AC) - nifHR (TTG TTS GCS GCR TAC ATS GCC ATC AT) for N-fixing bacterial DNA, amoA1F (GGG GHT TYT ACT GGT GGT) - amoA2R (CCC CTC KGS AAA GCC TTC TTC) for ammonia-oxidizing (nitrifying) bacterial DNA, nosZF (CAT GTG CAG NGC RTG GCA GAA) - nosZR (CGY TGT TCM TCG ACA GCC AG) for denitrifying bacterial DNA ^15^. The PCR conditions were: denaturation at 95°C for 10 min; 40 (for 16S rDNA), 45 (for ITS rDNA), or 50 (for others) cycles of 30 sec at 95°C, 45 sec at 55°C and 30 sec at 72°C; followed by dissociation curve analysis. Standard curves were calculated based on a serial dilution of gBlocks (linear DNA fragments) containing amplicon regions for each respective gene synthesized by Integrated DNA Technologies (Coralville, IA). All reactions were performed in triplicate in 384-well microplates using ABsolute qPCR Master Mix, containing SYBR Green and ROX, on an ABI 7900ht qPCR machine.

**Amplicon sequencing**

PCR amplification of both 16S and ITS amplicons from soil DNA extracts was carried out in 25 µl reactions including 1 µl of 20-fold diluted genomic DNA, 0.5 µl of each 10 µM primer, 5 µl of 10 × OneTaq Standard Reaction Buffer (New England BioLabs, Ipswitch MA), 0.5 µl of 10 mM dNTPs (New England BioLabs, Ipswitch MA), and 0.13 ul of Taq polymerase. All PCR reactions were set up on ice and using the hot start Taq polymerase (New England Biolabs, Ipswitch MA) to minimize non-specific amplification and primer dimerization. The PCR conditions for 16S were: denaturation at 94°C for 3 min; 35 cycles of 45 sec at 94°C, 1 min at 50°C and 1 min at 68°C; followed by a 5 min final extension at 68°C^16^. The PCR conditions for ITS were: denaturation at 94°C for 3 min; 35 cycles of 45 sec at 94°C, 1 min at 50°C and 1 min 30 sec at 72°C; followed by a 5 min final extension at 68°C^10,17^. The amplification was confirmed by electrophoresis. Amplicons were cleaned using Just-a-Plate 96 PCR Purification and Normalization Kit (Charm Biotech, MO, USA) and quantified using Qubit HS-dsDNA kit (Invitrogen, Carlsbad, California, USA) on a Tecan Infinite F200 Pro plate reader, reading at 485 nm excitation and 530 nm emission.

**Bioinformatics**

Sequencing resulted in a total of 18,740,961 16S sequences and 23,615,355 ITS sequences, with an average of 82,559 ± 33,915 and 104,032 ± 68,562 sequences per sample. The R package dada2 was used for sequence quality control, paired-end assembly, identification of amplicon sequence variants (ASVs), and taxonomy assignment^18^. Primers were removed by truncating the primer length bases and by using cutadapt^19^ for bacteria and fungi, respectively, and the reads were subject to quality filtering using the dada2 default setting. The filtered reads were then dereplicated and denoised, paired ends merged, and chimeric sequences removed. Taxonomy was assigned to ASVs using the naive Bayesian classifier method^20^ in combination with the UNITE database (v. 7.2)^21^ as the reference for fungal ITS ASVs and SILVA (release 138.1)^22^ as the reference for 16S ASVs. Mitochondria and chloroplasts were removed from the bacterial ASV table. The final, cleaned dataset consisted of a total of 12,419,929 16S sequences and 15,414,863 ITS sequences, with an average of 54,713 ± 26,521 16S sequences and 67,907 ± 40,312 ITS sequences per sample.

To assign fungal taxa to the functional guild, ASVs were searched against the FungalTrait database^23^. The FungalTrait database can assign lifestyle traits based on fungal genera. We selected ASVs with “ectomycorrhizal” and ‘saprotroph’ from the primary lifestyle columns. Saprotrophs were further categorized into the soil, wood, litter, dung, and unspecified saprotrophs. We also selected ASVs with any descriptions from the plant pathogenic capacity column and ASVs with the word “parasite” from the animal biotrophic capacity column. To search for the bacterial guild, bacterial genera were searched against a functional database created based on substrate enrichment experiments, and the presence of genes coding for enzymes in specific biochemical pathways (copiotroph, oligotroph, cellulolytic, ligninolytic, methanotroph, chitinolytic, assimilatory nitrite-reducing, dissimilatory nitrite-reducing, assimilatory nitrate-reducing, dissimilatory nitrate-reducing)^24,25^.

The “Suites” of bacteria and fungi were calculated for the urban edge, urban interior, rural edge, and rural interior location groups using the multipatt command in the indicspecies package^26^ in R. These suites were visualized in phylogenetic trees using MEGA X (Molecular Evolutionary Genetics Analysis) software^27^ and then illustrated using iTOL (Interactive Tree Of Life) software^28^. Since the ITS region is less suitable for phylogenetic analyses, 28S sequences downloaded from the SILVA ribosomal RNA database (version 138.1)^22^ were assigned for fungal ASVs^29^. The assignment was based on the genus name, but if the name was not found in the database, the family and order names were used instead. Any remaining ASVs were excluded from the analysis.

The PICRUSt2^30^ was used for predicting the bacterial xenobiotics degrading, plant pathogenic, and human disease infectious capacity, which were referred to in the KEGG pathway module “Xenobiotics biodegradation and metabolism”, the pathway “Plant-pathogen interactions”, and the pathway module “Infectious disease: bacterial”, respectively. The ASV table was normalized to the gene copy number, and metagenome functional profiles were predicted to generate a table of Kyoto Encyclopedia of Gene and Genomes (KEGG) Orthologues (KO)^31–33^.

**Network analysis**

To reduce noise and prevent spurious association predictions^34^, abundance tables were filtered to keep genera with at least 25 reads total over all network samples, a relative abundance greater than 0.1%, and present in more than 20% of network samples before networks were constructed. Abundance tables were additionally filtered to ensure that networks were not biased in other ways (i.e., community diversity, spatial autocorrelation of individual taxa). In *SpiecEasi*, we constructed networks using the sparse graphical lasso (glasso) setting and selected the optimal sparsity parameter based on the Stability Approach to Regularization Selection (StARS)^35^, with a variability threshold set to 0.1 for all networks. We calculated edge density (i.e., the number of connections in the network divided by the number of total possible connections if every genus was connected to every other genus) and transitivity (i.e., the tendency for genera to cluster in highly-connected groups) as measures of network connectivity. To evaluate the centrality, or importance, of genera, we calculated betweenness centrality (i.e., the proportion of shortest paths connecting two other genera in the network that pass through the genus) and degree centrality (i.e., the number of connections that a genus has) for each genus.

To construct synthetic metagenomes for bacterial and fungal networks, copy numbers of gene families were obtained for individual taxa from online databases. For bacterial taxa, the Metagenomics Inference PICRUSt2 pipeline^30^ was used to output the KEGG Orthology (KO)^31–33^ metagenome predictions for each bacterial ASV. For fungal taxa, Gene Ontology (GO)^36,37^ functional annotations were downloaded from the Joint Genome Institute MycoCosm^38^ for any genome within a genus contained in the dataset. Copy numbers of each bacterial KO number and fungal GO term were averaged within each genus and then weighted by the genus abundances in each network. Using the multipatt function in the indicspecies package of R^26^, bacterial KO numbers and fungal GO terms significantly associated with urban and rural networks (p < 0.05) were identified. The KEGG mapper reconstruct^32^ and QuickGO^39^ were used to determine the bacterial and fungal genetic functions associated with the KO numbers and GO terms, respectively. The Revigo^40^ interactive graph function was used to cluster GO terms into parent terms based on the 2023-01-01 release of the Gene Ontology database.

**Table S1** All eight study sites and their distance to Park Street Station in Boston, their % ISA, urbanization designation for each site, and additional topographic and tree community information. The two most common tree species at each site are shown in order of abundance. Harvard Forest sites are numbered 1–3 for clarity, differing from the numbering scheme in Garvey et al. (2022)^6^, but following Caron et al. (2023)^5^.

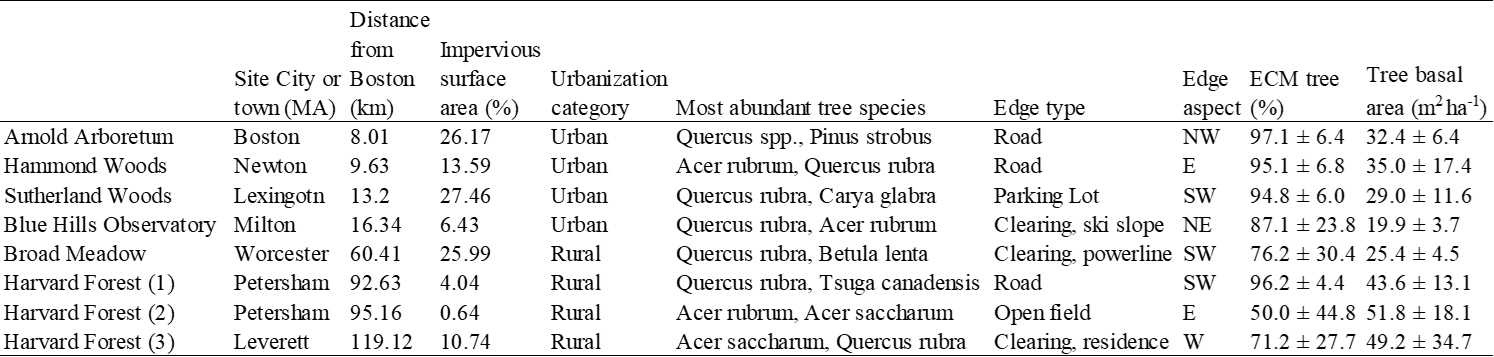

**Table S2** Urbanization and forest edge effects on (a) the abundance of fungal and bacterial groups. §, absolute abundance from qPCR (copies g^-1^ soil); ‡, the number of bacterial KEGG orthology counts predicted by PICRUSt2; No additional symbol, relative abundance from amplicon sequence (%). Because total tree basal area, ECM tree percentage, soil C and N contents, pH, and foliar C and N concentrations were not measured in all sampling years, only one- or two-year values of these factors were included. Model fixed effects include urbanization (negative distance from Boston), edge effects (negative distance from edge), and their interaction. Forest sites were included as a random effect in the model. An estimate of the correlation coefficient (*R*) from the linear mixed model and *F* values from the ANOVA test for the model are shown. qPCR data were log-transformed and all the data were standardized using the scale function of R before testing. Values with significant relationships to urbanization and forest edge are bolded. Asterisks represent *P* values (**P* < 0.05, ***P* < 0.01, ****P* < 0.001).

(a)

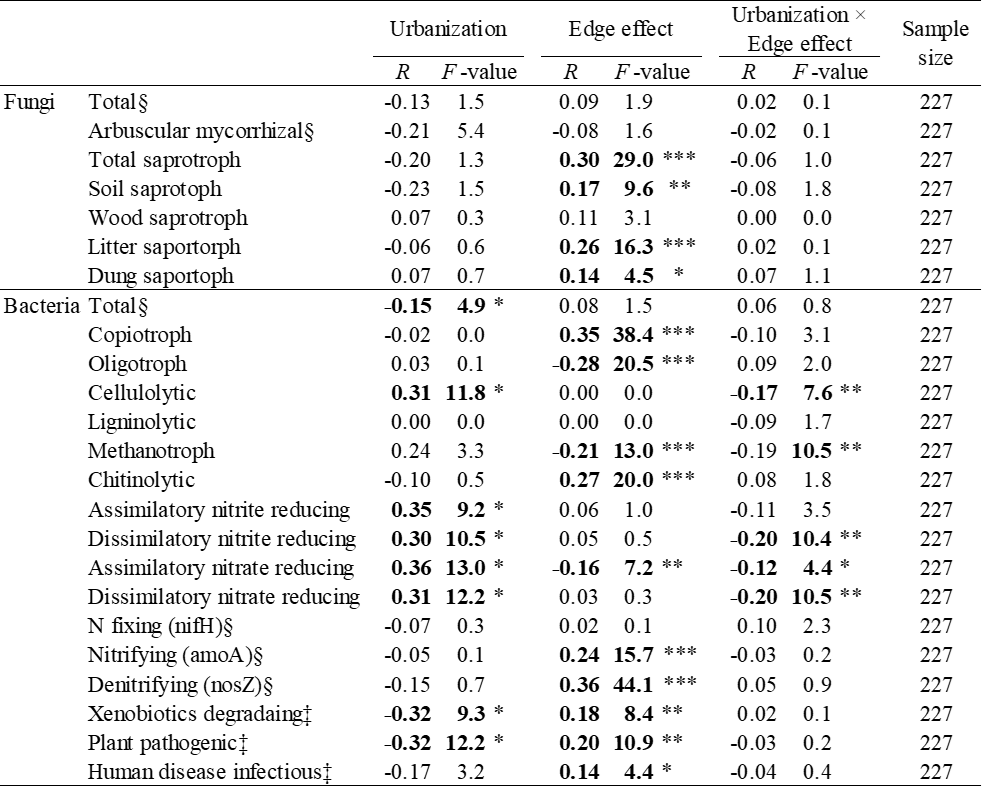

(b)

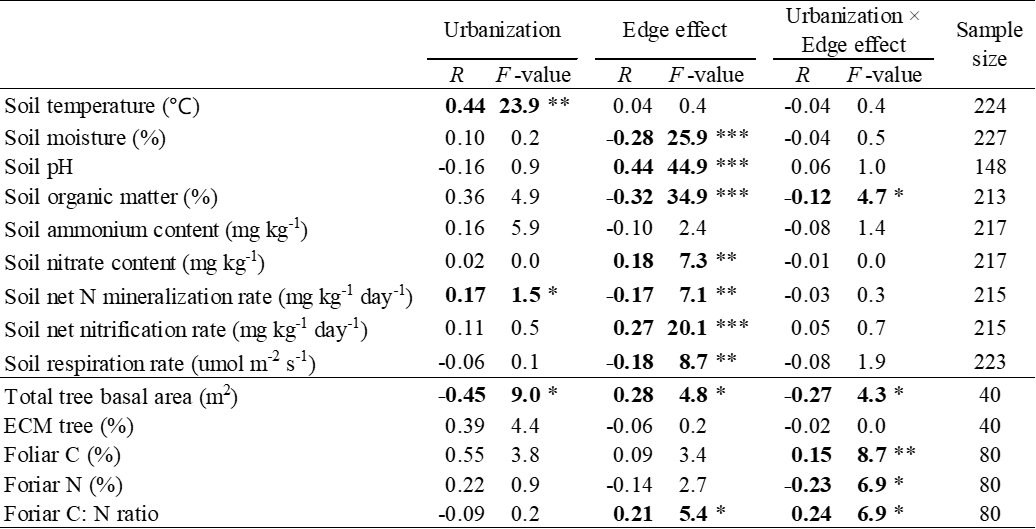

**Table S3** Urbanization and forest edge effects on the relative abundance of functional groups normalized by ANCOM-BC instead of the rarefication. Model fixed effects include urbanization (negative distance from Boston), edge effects (negative distance from edge), and their interaction. Forest sites were included as a random effect in the model. An estimate of the correlation coefficient (*R*) from the linear mixed model and *F* values from the ANOVA test for the model are shown. All the data were standardized using the scale function of R before testing. Values with significant relationships to urbanization and forest edge are bolded. Asterisks represent *P* values (**P* < 0.05, ***P* < 0.01, ****P* < 0.001).

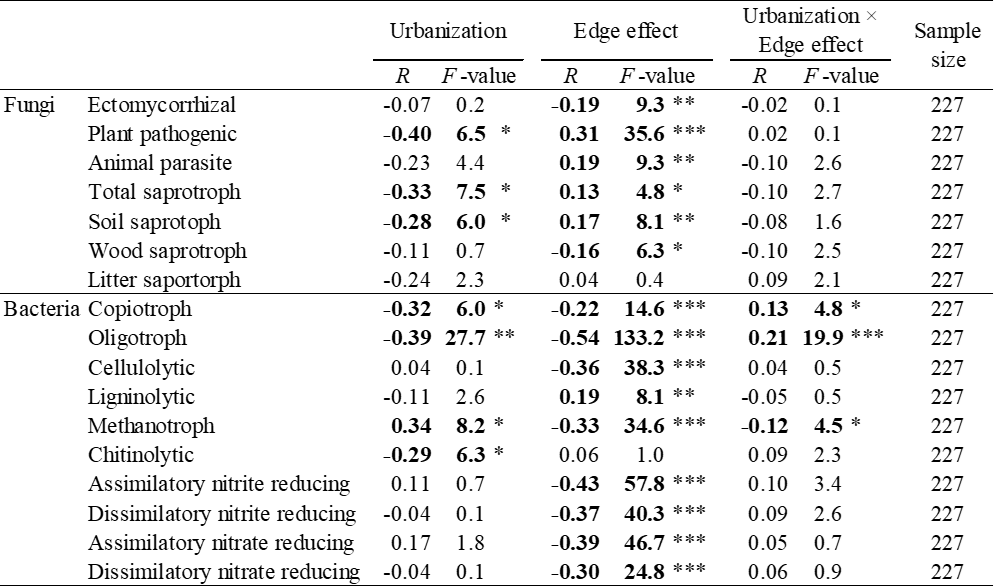

**Fig. S1**

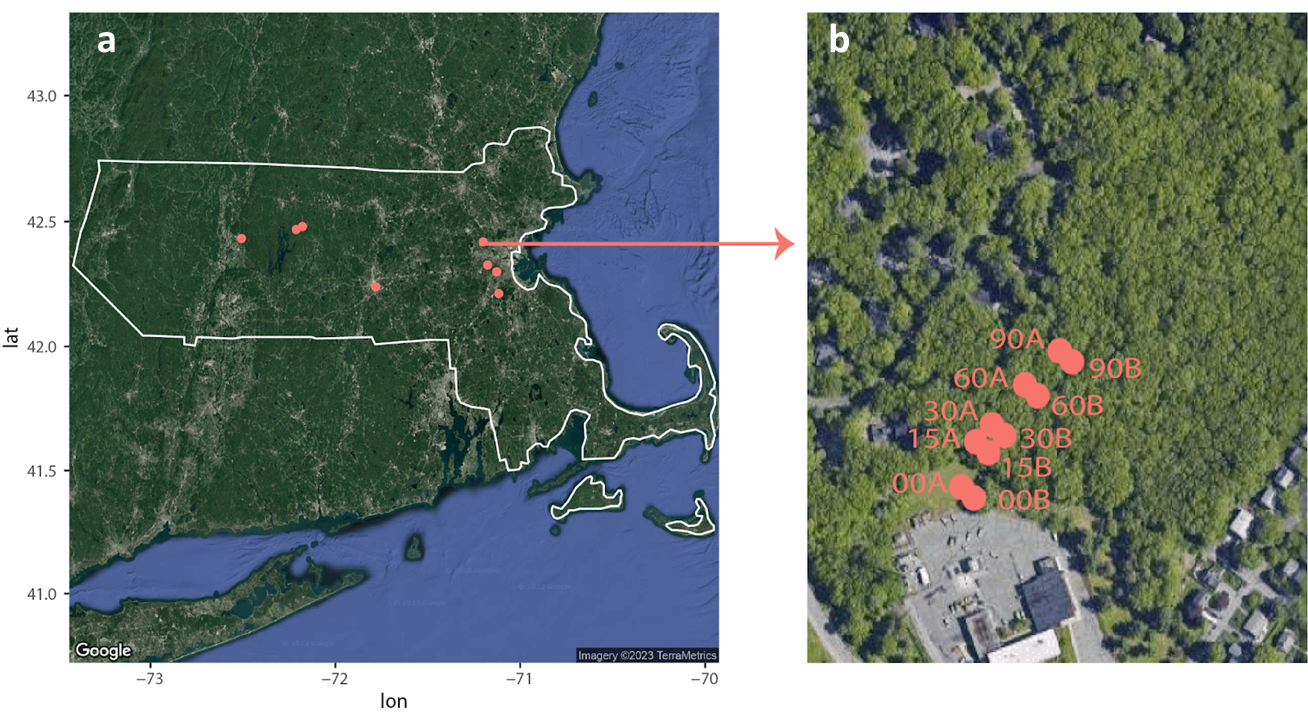

Fig. S1. Map of sampling regime and sampling locations across the UNE study system. Circles represent individual sampling locations mapped on Google Map images. (a) Individual sampling sites are shown on a map of the state of Massachusetts. (b) Individual sampling locations (replicates A and B) within a sampling site (Sutherland Woods) along the 0-90m distance from forest edge transect within an individual 100-m × 100-m plot are shown. lat, latitude; lon, longitude.

**Fig. S2**

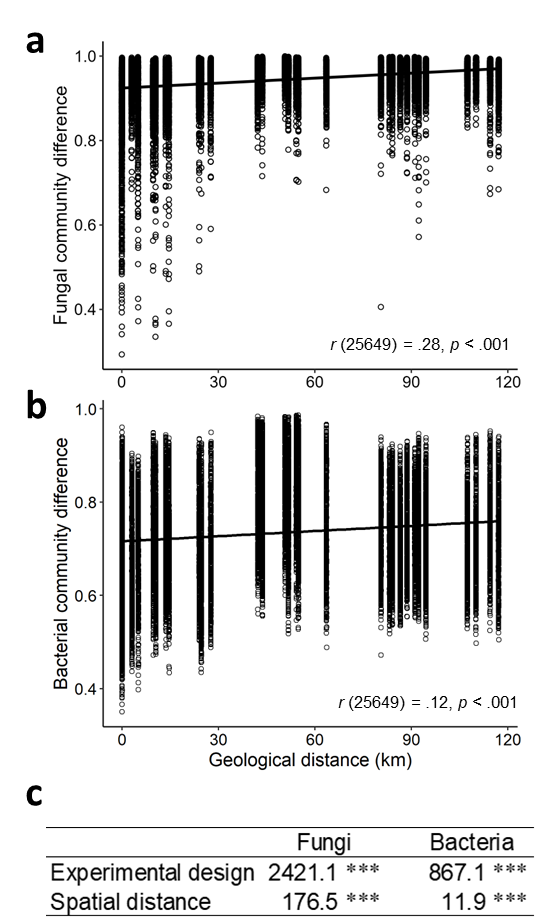

**Fig. S2** Mantel test between the (a) fungal and (b) bacterial community difference (Bray–Curtis dissimilarities) and the geological distance. (c) The result of Multiple Regression on distance Matrices (MRM) to test the effect of experimental design and spatial distance on fungal and bacterial communities. *F* values are shown, and asterisks represent *P* values (****P* < 0.001).

**Fig. S3**

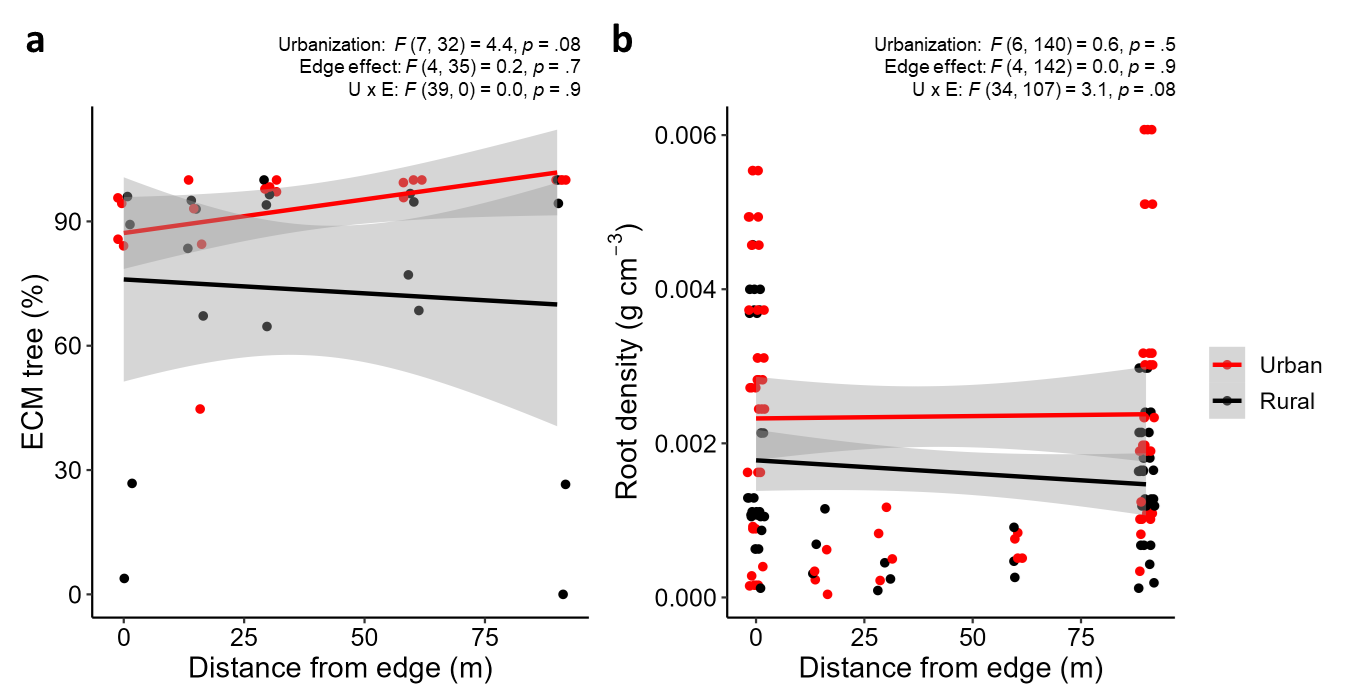

**Fig. S3** The relationship between distance from the edge and (a) ECM tree percentage based on basal area, and (b) the root density. The upper left shows the result of ANOVA for the linear mixed-effect model including urbanization (negative distance from Boston), edge effect (negative distance from edge), and their interactions (U×E) as the independent variables. Asterisks represent *P* values (**P* < 0.05).

**Fig. S4**

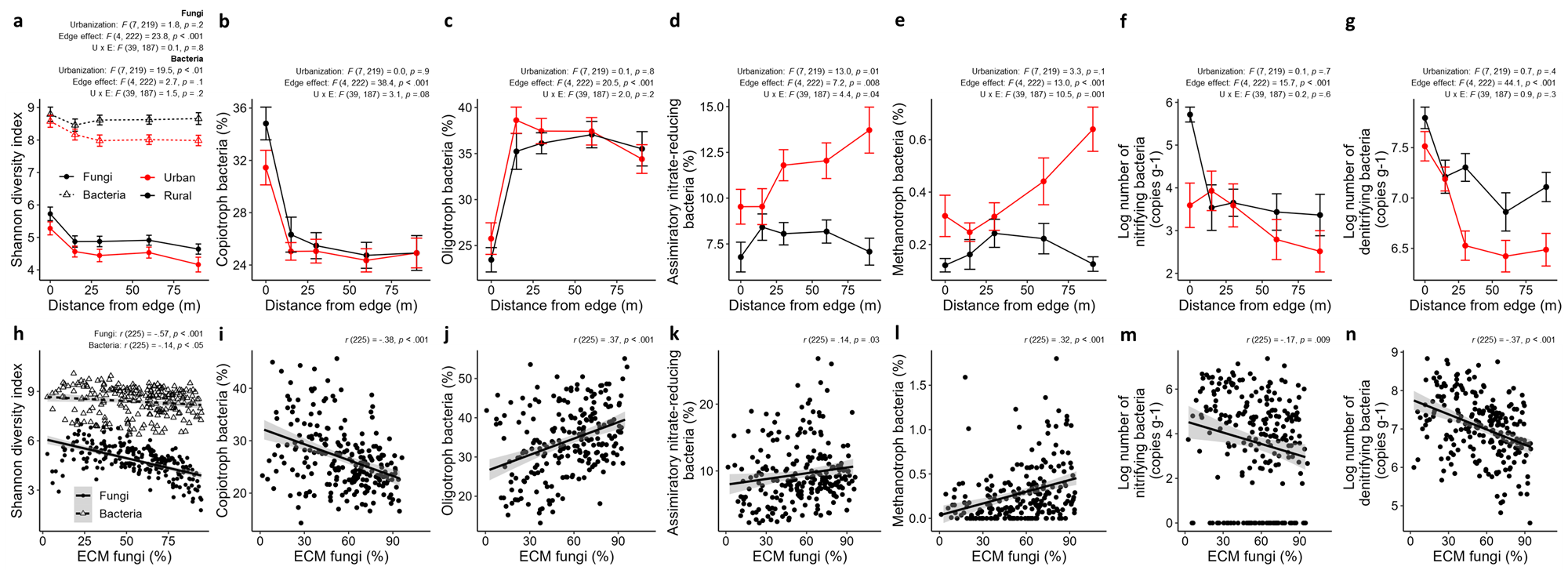

**Fig. S4** The relationship between distance from the edge and fungal/bacterial diversity or functional group abundance (a-g). Also shown is the relationship between ECM fungal relative abundance and fungal/bacterial diversity or functional group abundance (h-n). The upper left shows the result of ANOVA for the linear mixed-effect model for urbanization (negative distance from Boston), the edge effect (negative distance from edge), and their interaction (U×E) (a-g), or the correlation coefficient (h-n). Asterisks represent *P* values (**P* < 0.05, ***P* < 0.01, ****P* < 0.001).

**Fig. S5**

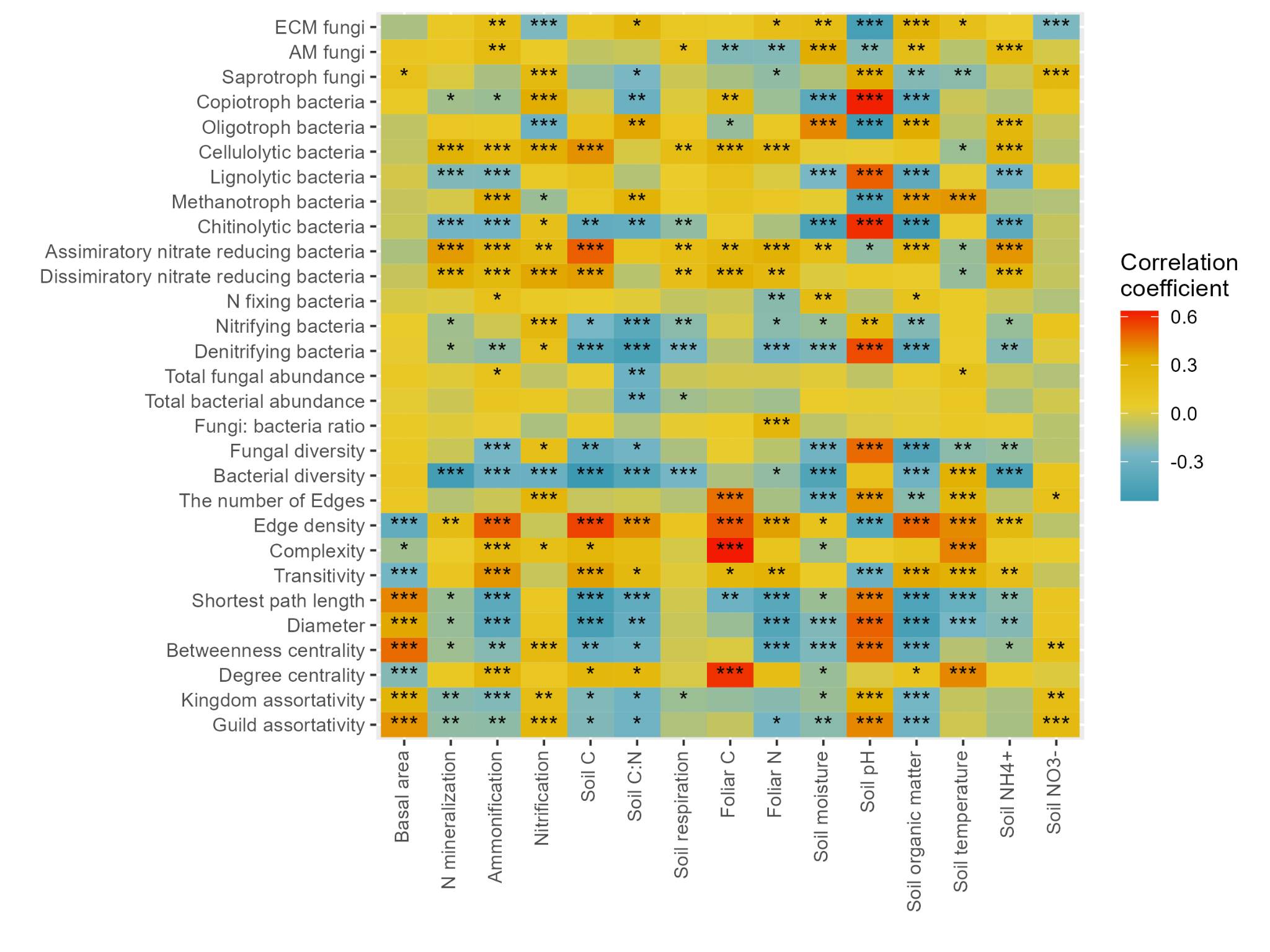

**Fig. S5** Correlation between microbial properties, biogeochemical pools and fluxes, and soil properties. The 10 rows from the bottom show the network parameters. *Pearson* correlation coefficient values are shown as colors. Asterisks represent *P* values (**P* < 0.05, ***P* < 0.01, ****P* < 0.001).

**Fig. S6**

N fertilization: *F* (2,6) = 13.0, *p* < .01

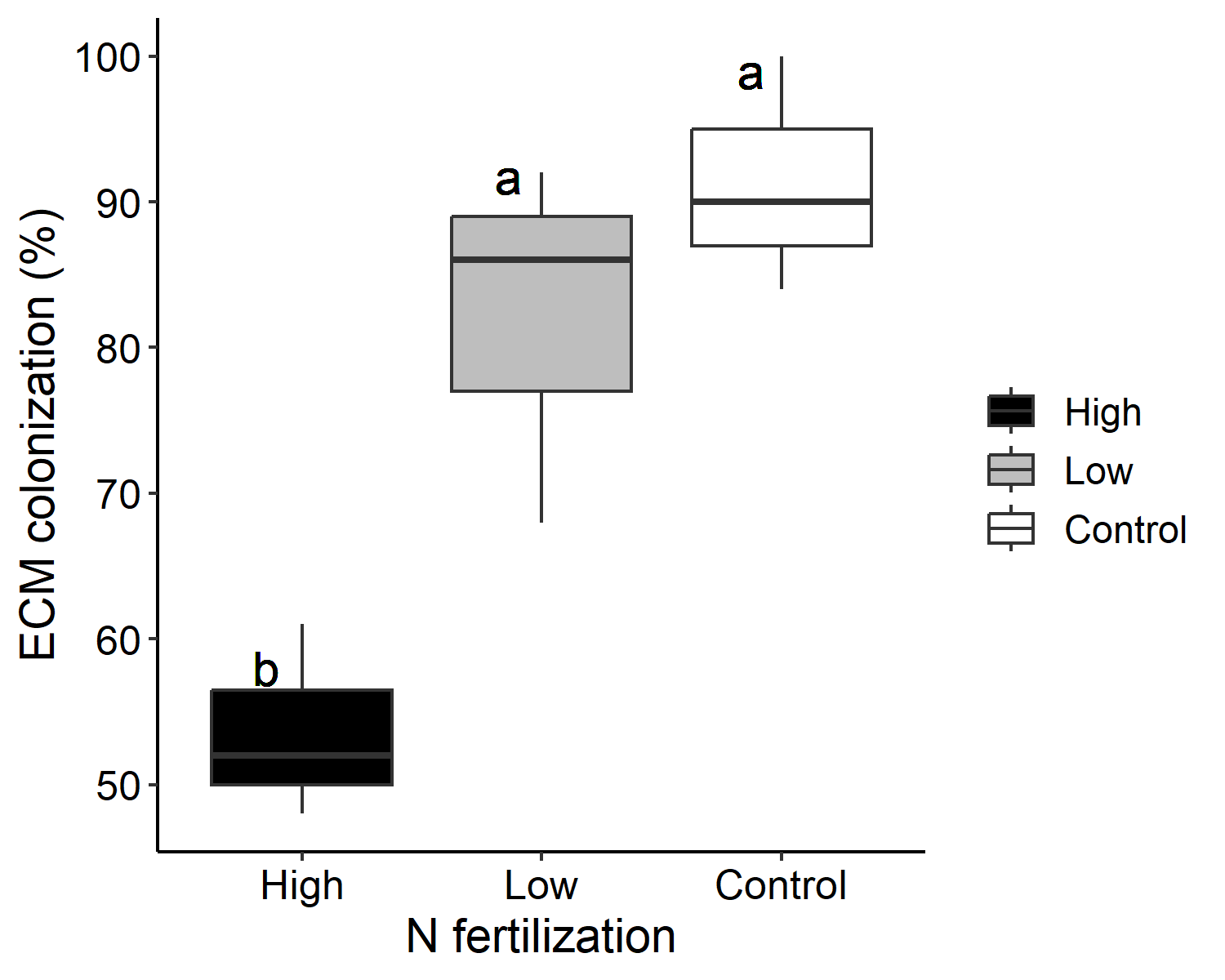

**Fig. S6** ECM colonization of oak tree roots from the Chronic N Amendment Experiment at Harvard Forest^8^. High and low N addition plots have received 150 kg N ha^-1^ yr^-1^1 and 50 kg N ha^-1^ yr^-1^, (respectively, every year from 1988 to 2019 year. N is applied as NH_4_NO_3._ Statistical results of ANOVA for the linear model testing for effects of fertilization level are shown. Different alphabet letters show significant differences based on Tukey's multiple comparison tests.

**Fig. S7**

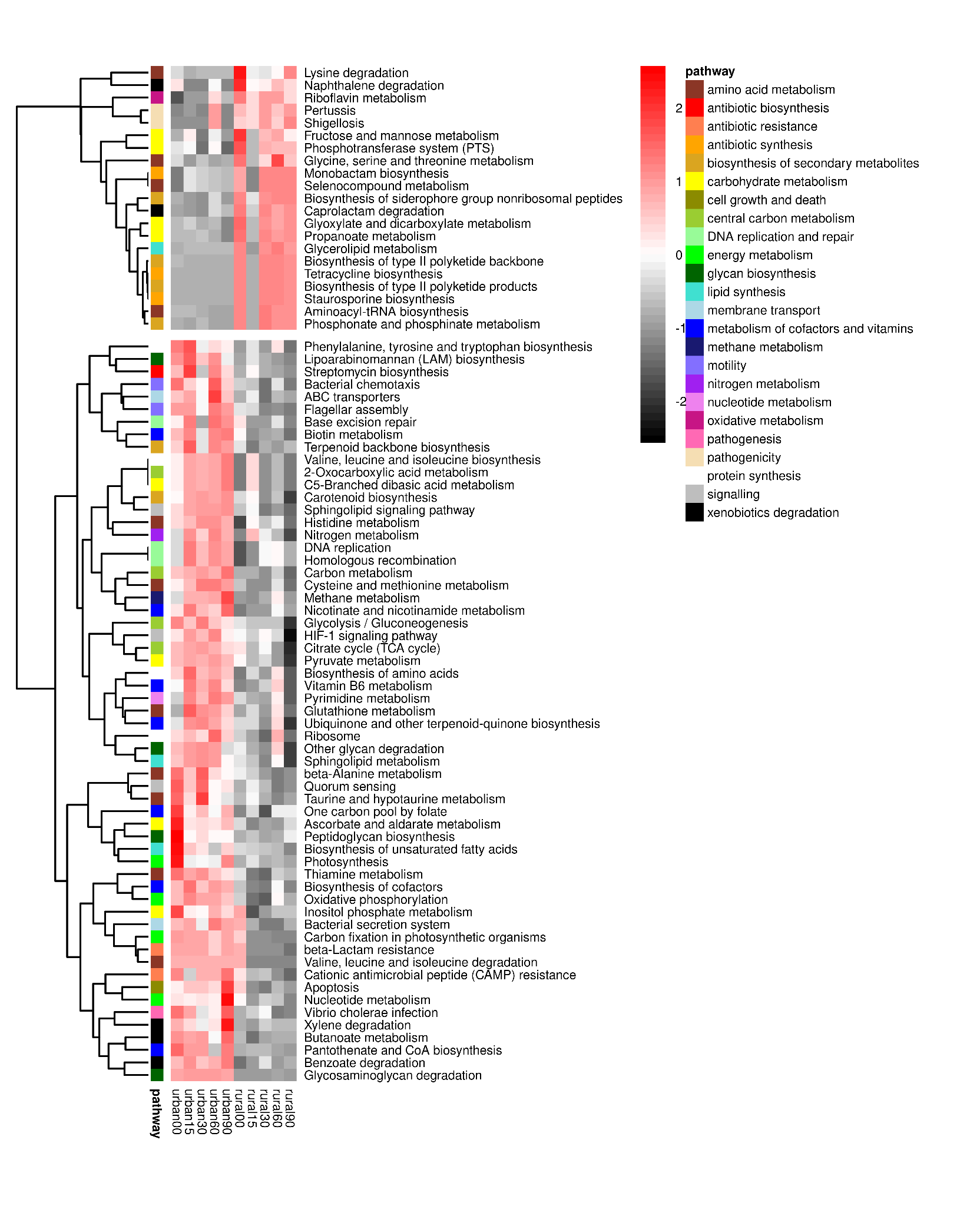

**Fig. S7** Heatmaps show the bacterial gene groups enriched in urban and rural networks, expanding upon the pathways from Fig. 2e. Gene group abundances are scaled to z-scores by row in the heatmap. Gene groups above the heatmap break are enriched in rural networks, while gene groups below the heatmap break are enriched in urban networks.

**Fig. S8**

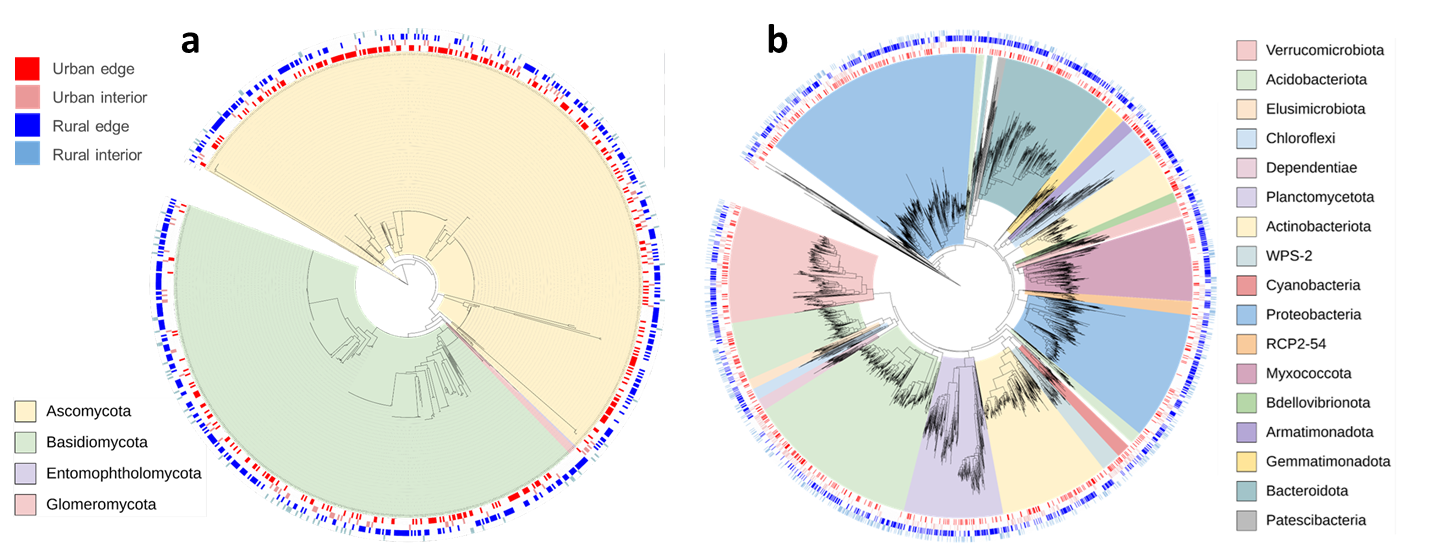

**Fig. S8** Phylogenetic trees of (a) fungal and (b) bacterial ASV detected as the indicator species in each forest type (Urban edge, Urban interior, Rural edge, and Rural interior). Phyla are indicated by different colors on the interior of the tree, while the forest-type designation is indicated by the outer rings.

**Fig. S9**

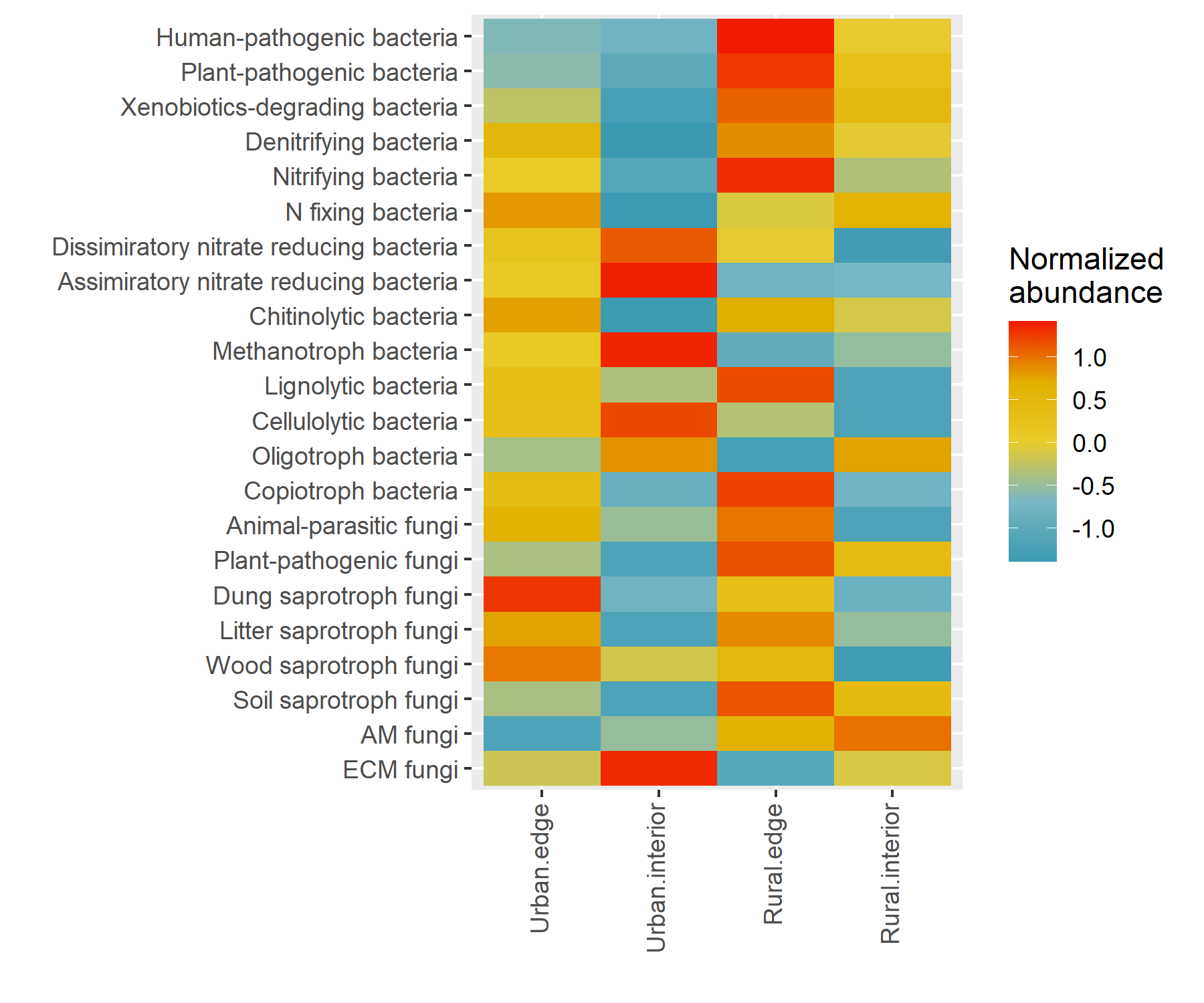

**Fig. S9** The relative abundance of microbial functional groups in each forest type (Urban edge, Urban interior, Rural edge, and Rural interior). The units used for each abundance are the same as in Table S2. The abundance was normalized in each row using the scale function in R.

**Fig. S10**

(a)	(b)

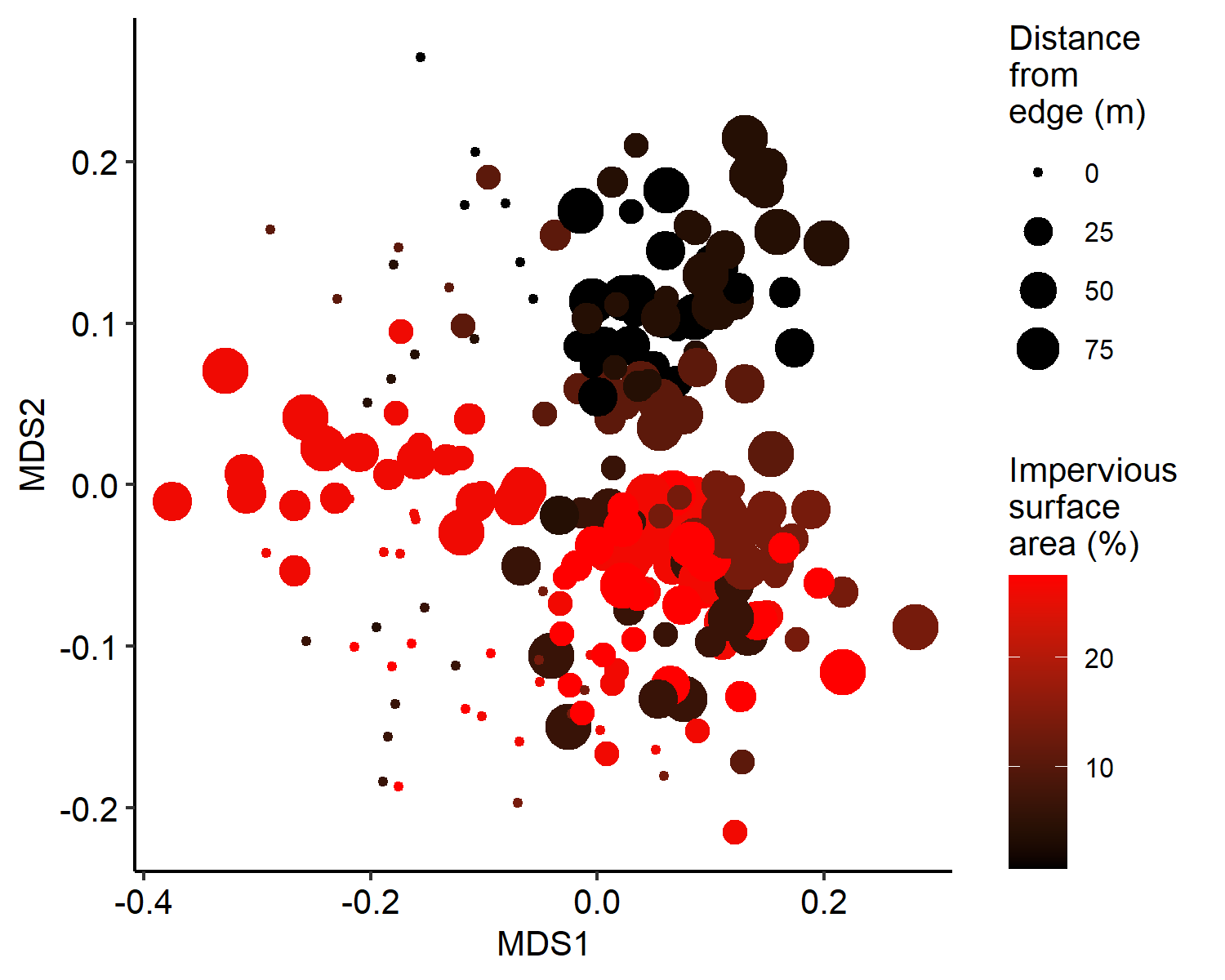

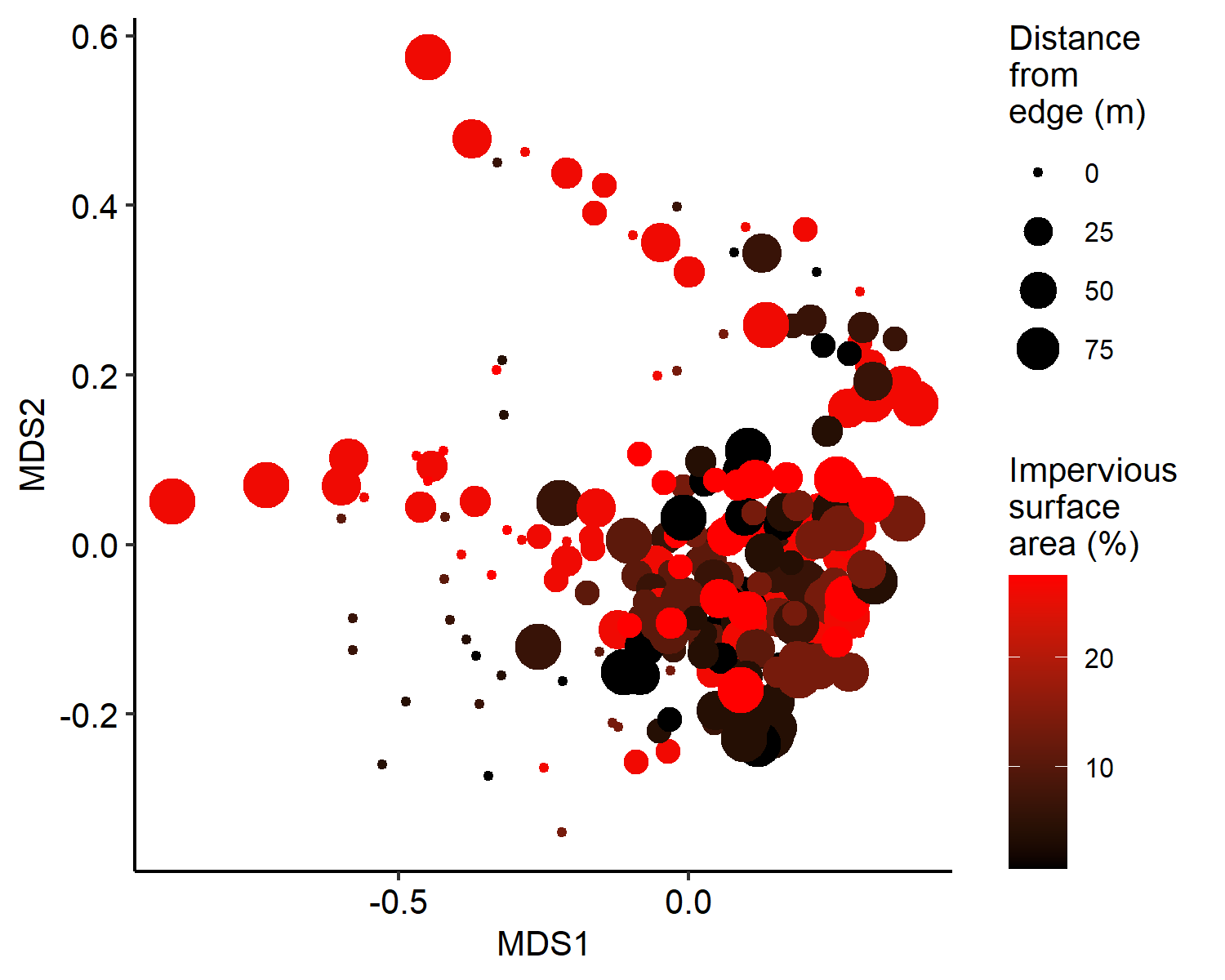

**Fig. S10** Nonmetric multidimensional scaling (NMDS) based on Bray–Curtis dissimilarities of (a) fungal and (b) bacterial community composition. The color and size of the points represent the percent impervious surface area (ISA) and distance from the edge (m), respectively.
